# Baricitinib treatment resolves lower airway inflammation and neutrophil recruitment in SARS-CoV-2-infected rhesus macaques

**DOI:** 10.1101/2020.09.16.300277

**Authors:** Timothy N. Hoang, Maria Pino, Arun K. Boddapati, Elise G. Viox, Carly E. Starke, Amit A. Upadhyay, Sanjeev Gumber, Kathleen Busman-Sahay, Zachary Strongin, Justin L. Harper, Gregory K. Tharp, Kathryn L. Pellegrini, Shannon Kirejczyk, Keivan Zandi, Sijia Tao, Tristan R. Horton, Elizabeth N. Beagle, Ernestine A. Mahar, Michelle YH Lee, Joyce Cohen, Sherrie M. Jean, Jennifer S. Wood, Fawn Connor-Stroud, Rachelle L. Stammen, Olivia M. Delmas, Shelly Wang, Kimberly A. Cooney, Michael N. Sayegh, Lanfang Wang, Daniela Weiskopf, Peter D. Filev, Jesse Waggoner, Anne Piantadosi, Sudhir P. Kasturi, Hilmi Al- Shakhshir, Susan P. Ribeiro, Rafick P. Sekaly, Rebecca D. Levit, Jacob D. Estes, Thomas H. Vanderford, Raymond F. Schinazi, Steven E. Bosinger, Mirko Paiardini

**Affiliations:** Division of Microbiology and Immunology, Yerkes National Primate Research Center, Emory University, Atlanta, Georgia, USA; Yerkes Genomics Core Laboratory, Yerkes National Primate Research Center, Emory University, Atlanta, Georgia, USA; Division of Animal Resources, Yerkes National Primate Research Center, Emory University, Atlanta, Georgia, USA; Department of Medicine, School of Medicine, Emory University, Atlanta, Georgia, USA; Department of Radiology and Imaging Sciences, School of Medicine, Emory University, Atlanta, Georgia, USA; Department of Pathology and Laboratory Medicine, School of Medicine, Emory University, Atlanta, Georgia, USA; Division of Pathology, Yerkes National Primate Research Center, Emory University, Atlanta, Georgia, USA; Center for AIDS Research, Department of Pediatrics, School of Medicine, Emory University, Atlanta, Georgia, USA; Vaccine and Gene Therapy Institute, Oregon Health & Science University, Beaverton, OR, USA; Oregon National Primate Research Center, Oregon Health & Science University, Beaverton, OR, USA; Department of Pathology, Case Western Reserve University, Cleveland, OH, USA; Center for Infectious Disease and Vaccine Research, La Jolla Institute for Immunology, La Jolla, CA, USA

**Keywords:** COVID-19, inflammation, SARS-CoV-2, immune activation, baricitinib, nonhuman primate, immunology, pharmacology, pathogenesis

## Abstract

Effective therapeutics aimed at mitigating COVID-19 symptoms are urgently needed. SARS-CoV-2 induced hypercytokinemia and systemic inflammation are associated with disease severity. Baricitinib, a clinically approved JAK1/2 inhibitor with potent anti-inflammatory properties is currently being investigated in COVID-19 human clinical trials. Recent reports suggest that baricitinib may also have antiviral activity in limiting viral endocytosis. Here, we investigated the immunologic and virologic efficacy of baricitinib in a rhesus macaque model of SARS-CoV-2 infection. Viral shedding measured from nasal and throat swabs, bronchoalveolar lavages and tissues was not reduced with baricitinib. Type I IFN antiviral responses and SARS-CoV-2 specific T cell responses remained similar between the two groups. Importantly, however, animals treated with baricitinib showed reduced immune activation, decreased infiltration of neutrophils into the lung, reduced NETosis activity, and more limited lung pathology. Moreover, baricitinib treated animals had a rapid and remarkably potent suppression of alveolar macrophage derived production of cytokines and chemokines responsible for inflammation and neutrophil recruitment. These data support a beneficial role for, and elucidate the immunological mechanisms underlying, the use of baricitinib as a frontline treatment for severe inflammation induced by SARS-CoV-2 infection.

## Main

The rapid emergence and dissemination of severe acute respiratory syndrome coronavirus 2 (SARS-CoV-2) and subsequent COVID-19 pandemic has placed an excessive burden on public and private healthcare systems with over 800,000 deaths worldwide. Thus, therapeutic approaches aimed at mitigating disease severity are of utmost global priority (https://www.who.int/). SARS-CoV-2 is highly transmissible and results in a wide spectrum of disease severity, ranging from asymptomatic individuals to critically-ill patients leading to death. Severe COVID-19 disease presents with high-grade fever, dry cough, pneumonia, bilateral opacities seen in radiographs, inflammation of the lungs and infiltration of immune cells. It has been noted that individuals with co-morbidities and compromised immune systems are at higher risk for severe clinical manifestations (Guan et al., 2020).

Immunological features of COVID-19 progression includes a robust pro-inflammatory response driven by innate and adaptive immune cells. Severe cases of COVID-19 have been shown to have elevated serum levels of pro-inflammatory cytokines and chemokines including: IFNγ, TNFα, IP-10, G-CSF, IL-2, IL-6 IL-8, IL-9, IL-10, and IL-17 (Huang et al., 2020). Therefore, the use of therapeutics targeted at Janus Kinases have the potential to ameliorate disease severity by limiting the hypercytokinemia and cytokine release syndrome (CRS) seen in COVID-19 patients (Huang et al., 2020).

Non-human primate (NHP) models have been used extensively to study pathogenesis and potential vaccine and antiviral candidates for numerous viral diseases (Estes et al., 2018). We and others have recently used rhesus macaques (RMs) to model SARS-CoV-2 infection and pathogenesis; SARS-CoV-2 infected RMs develop transient respiratory disease and exhibit viral shedding similar to humans, recapitulating mild to moderate infection, and in rare cases severe disease (Chandrashekar et al., 2020b; Munster et al., 2020; Rockx et al., 2020; Williamson et al., 2020; Yu et al., 2020). Here we use RMs to assess baricitinib monotherapy as a candidate therapeutic for the treatment of COVID-19. Baricitinib is an oral, selective inhibitor of Janus Kinases (JAK1/2) with potent anti-inflammatory activity approved for treatment of patients with moderate to severe active rheumatoid arthritis (Keystone et al., 2015). Recently, machine learning algorithms and *in vitro* data suggested that baricitinib could also inhibit clathrin-mediated endocytosis of SARS-CoV-2 (Cantini et al., 2020; Richardson et al., 2020; Stebbing et al., 2020; Titanji et al., 2020). Thus, baricitinib could provide a dual effect of dampening inflammation and reducing viral replication. In this study, leveraging the ability to perform longitudinal collections, including bronchoalveolar lavages, and the availability of lung tissue for pathology, we tested the immunologic and virologic effects of baricitinib treatment in SARS-CoV-2 infected RMs.

## Results

### Baricitinib was well-tolerated and detectable in plasma and tissues, but did not limit viral replication in SARS-CoV-2 infected RMs

We inoculated 8 adult RMs (11-17 years old, mean = 14 years, **supplementary table 1**) with a total of 1.1x10^6^ PFU SARS-CoV-2 (2019-nCoV/USA-WA1/2020), administered by intranasal (IN) and intratracheal (IT) routes(Yu et al., 2020). Two days post viral inoculation, 8 RMs were randomized to receive 4 mg of oral baricitinib, daily for 8-9 days or observed without treatment until day 10-11 post infection (DPI) when all RMs were euthanized (**Fig.1a**). We first determined the concentration and biodistribution of baricitinib in the treated group. At 24 hours post dose, baricitinib was readily detected in plasma of all treated animals (**Fig.1b**; measures performed at 6 DPI closed symbol; and 8 DPI open symbol), achieving an average level of 2.13 ng/mL. At necropsy, baricitinib was detectable at approximately 2 hours after the last dose in left/right upper and lower lung (**Fig.1c;** n = 4 RMs; average of 4.41 and 4.43 ng/g, respectively), brain (n = 3 RMs; 2.09 ng/g tissue) and cerebrospinal fluid (CSF; n = 2 RMs; 0.29 ng/ml) (**Fig. S1**); we also detected baricitinib in CSF from 3 out of the 4 treated animals at 24 hours post dosing on 8-9 days after the final dose (**Fig. S1**).

**Figure 1.**
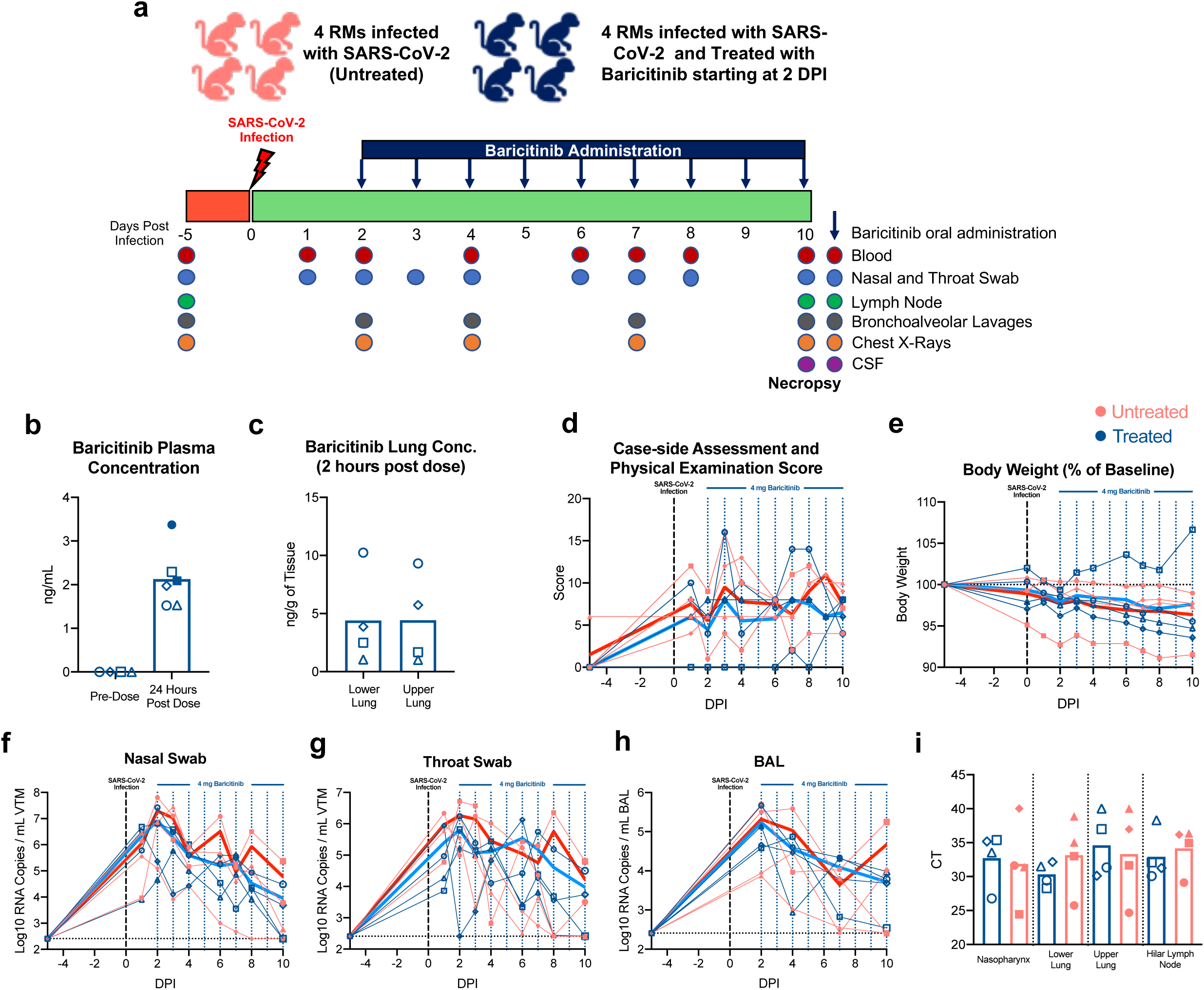
Baricitinib is detectable in plasma and tissues from SARS-CoV-2 infected RMs, but has not impact viral kinetics. (**a**) Study design; 8 RMs were infected intranasally and intratracheally with SARS-CoV-2 and at 2 days post-infection (DPI), 4 RMs began daily baricitinib administration (4 mg). Longitudinal collections performed are indicated in circles. (**b**) Concentration of baricitinib 24 hours post-dosing in plasma (6 DPI closed symbol; and 8 DPI open symbol) and (**c**) at necropsy in upper and lower lungs of baricitinib treated SARS-CoV-2 infected RMs. (**d**) Daily cage-side assessment and physical examination scores and (**e**) changes in body weight from baseline in baricitinib treated (blue symbols; n = 4) and untreated (red symbols; n = 4) SARS-CoV-2 infected RMs. (**f-g**) After SARS-CoV-2 inoculation, nose and throat swabs, and bronchoalveolar lavages (BAL) were collected and viral loads were quantified by qRT-PCR. (**i**) Viral loads in tissues measured at necropsy (10-11 DPI). Ct, cycle threshold. Different symbols represent individual animals. Thick lines represent the average of the baricitinib treated (blue lines), and untreated groups (red lines). Bars in b, c, and j, represent the average of the treated and untreated groups. Statistical analysis was performed using a non-parametric Mann-Whitney Test.

In terms of potential side effects, a slight reduction of peripheral monocytes, neutrophils and lymphocytes, which could be due to trafficking to the lung, as well as decreased red blood cell counts (RBC), hematocrit (HCT) and hemoglobin (HGB) were observed starting at 2 DPI in all RMs (**Fig. S2 a-f**). Blood chemistries showed elevated levels of alkaline phosphatase (ALP) in one untreated animal starting at 2 DPI, and all other values were within the normal range (**Fig. S2 g**). Thus, treatment with baricitinib was well-tolerated without direct evidence of treatment-induced clinical pathology, abnormal blood counts, nephrotoxicity or hepatotoxicity when compared to untreated SARS-CoV-2 infected animals.

To further monitor response to infection and baricitinib treatment, the health status of all animals was assessed daily by veterinarians, with cage-side assessment and physical examination scored based on a standardized scoring system (modified from previous studies (Chertow et al., 2016; Huang et al., 2020); main parameters included in the scoring are listed on **Supplementary Table 2 and 3**). On 1 DPI, all animals exhibited changes to alertness and respiratory pattern (**Fig.1d**). Additional early signs of disease included: changes to pulse oximetry readings, reduction in appetite, hunched posture, shivering, pale appearance and agitation. Signs of disease persisted during the 10/11-day course of the study, without differences between treated and untreated animals (**Fig.1d**). Weight loss was observed in 4/4 untreated and 3/4 baricitinib treated animals (**Fig.1e**), although we cannot discriminate if this is a result of the infection or related to frequent access for sample collection.

We next assessed viral RNA levels by qRT-PCR in multiple anatomic sites (Chandrashekar et al., 2020b; Munster et al., 2020). We observed high levels of SARS-CoV-2 RNA in nasal and throat swabs, and bronchoalveolar lavages (BAL), with a peak between days 2-4 post-infection of **1.4x10^7^, 1.2x10^6^, and 1.9x10^5^ copies/mL** respectively (**Fig.1f-h**); viral RNA then steadily decreased until 10-11 DPI. SARS-CoV-2 RNA levels remained similar in nasal, throat, or BAL between the baricitinib treated and the untreated group. Virus was not detected in blood. At necropsy (10-11 DPI), viral RNA was detected for all animals in nasopharynx, lower and upper lungs, and hilar lymph nodes. Viral loads (cycle threshold value) in tissue for treated and untreated animals were comparable (**Fig.1i**). Additionally, *in situ* RNA hybridization (RNAscope) targeting both positive and negative-sense (data not shown) viral RNA strands identified multifocal clusters of infected cells within the lung parenchyma in both treated and untreated animals (**Fig. S3**). Thus, baricitinib treatment starting at 2 DPI was safe and well tolerated, but did not impact the kinetics of SARS-CoV-2 replication.

### Baricitinib reduced lung pathology and inflammation in SARS-CoV-2 infected rhesus macaques

We then performed multiple analyses to determine the severity of SARS-CoV-2 infection in RMs and the effectiveness of baricitinib to ameliorate the pathophysiologic response. First, x-ray radiographs (RM6 x-ray; **Fig.2a)** were longitudinally (-5, 2, 4, 7 and 10 DPI) performed (blinded scoring by a radiologist as previously reported(Munster et al., 2020; Williamson et al., 2020)). Pulmonary infiltration and ground glass opacity were observed at multiple experimental timepoints PI in 2/4 untreated and 0/4 treated RMs (**Fig.2b and c**), with one of the untreated animals showing severe pneumonia at all PI time points (**Fig. 2b and c**). Second, we measured serum levels of several systemic inflammatory markers. Among these, ferritin (4 DPI, p=0.0286; and 10 DPI, p=0.0286) and C-reactive protein (CRP; 4 DPI) levels were found to be elevated in the untreated RMs when compared with the baricitinib treated RMs (**Fig.2d and e**). Previous reports have indicated that heightened ferritin and CRP levels are indicative of COVID-19 severity in humans (Qin et al., 2020; Ruan et al., 2020). Finally, to assess lung damage of SARS-CoV-2 infection, all RMs were euthanized at 10 or 11 DPI. At necropsy, multiple regions of upper, middle and lower lung lobes were taken for immunologic, virologic and pathologic analyses. Lung pathologic analyses and scoring were performed by two pathologists independently in a blinded fashion. Treated animals showed decreased type 2 pneumocyte hyperplasia, peribronchiolar hyperplasia, syncytia formation, alveolar septal thickening and inflammatory cell infiltration (**Fig.2f-k**). Consistent with the pathology scoring, neutrophil (myeloperoxidase, MPO^+^, cells) and macrophage (ionized calcium-binding adaptor molecule, Iba-1^+^, cells) infiltration appeared to be decreased in the lungs in baricitinib treated animals as measured by quantitative immunohistochemistry (IHC) (**Fig. S4**). Of note, some of the SARS-CoV-2 infected animals, in both groups showed cell infiltration levels similar to uninfected RMs, indicating a resolution of the infiltration at 10 DPI, consistent with an earlier peak of pathogenesis in RMs, as previously published (Chandrashekar et al., 2020b; Munster et al., 2020; Rockx et al., 2020; Williamson et al., 2020; Yu et al., 2020). The average pathology score per lobe (measuring the average severity of abnormalities per lobe, independently of how many lobes had been effected, p=0.0286) and the total pathology score (considering severity and number of effected lobes, p=0.0857) were lower in the baricitinib treated group (0.99 and 22, respectively) as compared to untreated RMs (1.66 and 38.5, respectively) (**Fig.2l and n**). Overall, these data support a therapeutic role of baricitinib in reducing lung pathology, infiltration of inflammatory cells in the lung, and soluble markers of inflammation associated with disease progression in humans.

**Figure 2.**
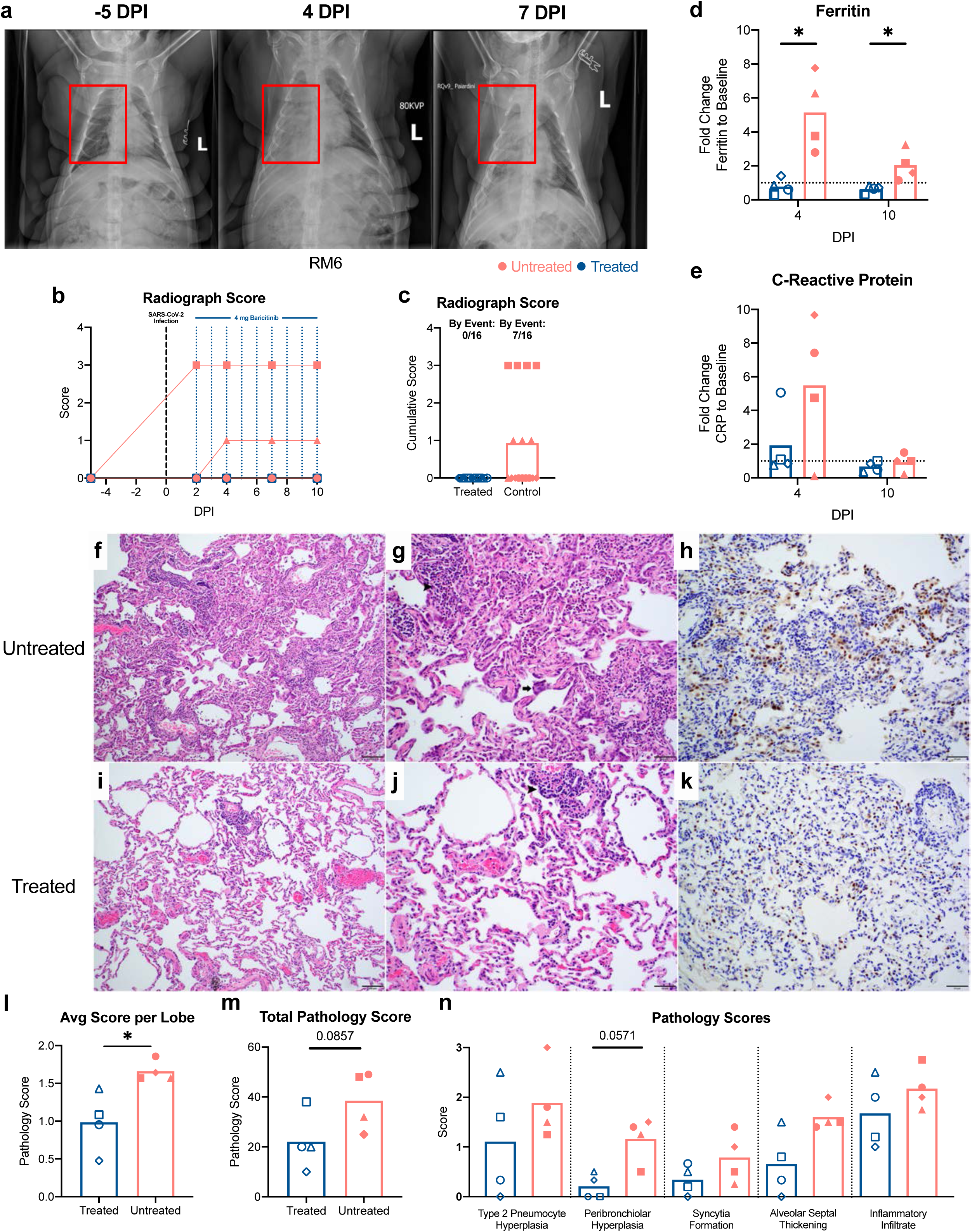
Reduced respiratory disease and lower levels of lung pathology in baricitinib treated RMs. (**a**) Representative ventrodorsal radiograph of an untreated RM before SARS-CoV-2 infection (-5 DPI), and at 4, and 7 DPI. Red squares indicate regions of pulmonary infiltrates and opacity. (**b**) Daily and (**c**) cumulative radiograph scores; ventrodorsal and lateral radiographs were scored for the presence of pulmonary infiltration by a clinical radiologist according to a standard scoring system (0: normal; 1: mild interstitial pulmonary infiltrates; 2: moderate pulmonary infiltrates with partial cardiac border effacement and small areas of pulmonary consolidation; 3: severe interstitial infiltrates, large areas of pulmonary consolidation, alveolar patterns and air bronchograms). Fold change of coagulation markers to 2 DPI for ferritin (**d**) and C reactive protein (CRP) levels (**e**). Panel (**f**) and (**g)** show representative lung lesions in an untreated SARS-Cov2 infected rhesus macaque with focally extensive interstitial pneumonia, type 2 pneumocytes hyperplasia, alveolar septal thickening, syncytia formation (arrow), neutrophils and macrophages infiltrations (arrowhead). Panel (**h**) shows Thyroid Transcription Factor-1 (TTF-1) staining with prominent type 2 pneumocyte hyperplasia (brown) in a control SARS-Cov2 infected rhesus macaque. Panel (**i**) and (**j)** shows treatment effects of baricitinib in COVID-19 infected rhesus macaques with a reduction in pulmonary lesions (affected area marked by arrows), lesser inflammatory infiltrates (arrowhead) and reduced type 2 pneumocyte hyperplasia. Panel (**k**) shows TTF-1 staining with lesser type 2 pneumocyte hyperplasia (brown) after Baricitinib treatment. (**l**) Average pathology score per lobe. (**m**) Total Pathology Score. (**n**) Pathology scores for individual parameters. Magnification, Panels (**g**) and (**i**): 100 ×; Panels (**g**), (**h**), (**j**), (**k**): 200 ×. Scale bar, panels (**f**) and (**i**): 200μM; Panels (**g**), (**h**), (**j**) and (**k**): 50μM. Pathology scoring. Bars in (**d**), (**e**), (**l)**, (**m**) and (**n**) indicate mean values for baricitinib treated (blue), and untreated (red) SARS-CoV-2 infected RMs. Each symbol represents individual animals. Statistical analysis in b, and j were performed using non-parametric Mann-Whitney Test.

### Baricitinib treatment dampens gene signatures of inflammation and neutrophil degranulation in the BAL of SARS-CoV-2 infected rhesus macaques

To investigate the impact of baricitinib on the lower airway, we performed bulk RNA-Seq profiling of cells isolated from BAL prior to SARS-CoV-2 inoculation (-5 DPI); 2 days after virus inoculation, prior to baricitinib treatment (2 DPI); and 4 days after infection, and 48 hours after beginning baricitinib (4 DPI). Relative to pre-infection, we observed a robust upregulation of differentially expressed genes (DEGs) at 2 DPI in both the treated and untreated animals (**Fig.3a)**, however at 4 DPI only a handful of DEGs were detected in the baricitinib treated animals, whereas a robust transcriptional response persisted in the untreated group. To identify immunological pathways perturbed by (i) SARS-CoV-2 infection, and (ii) baricitinib treatment, we performed gene-set enrichment analysis (GSEA)(Subramanian et al., 2005). To determine pathways that changed after drug administration, we directly compared gene expression profiles at 2 DPI to 4 DPI. Comparison of GSEA data from 2 DPI to 4 DPI in untreated animals show robust, highly significant positive enrichment in pathways comprised of genes for inflammatory responses, TNFα and IL6 signaling, neutrophil and granulocyte function – indicating that, in the absence of baricitinib, expression of these genes, continues to increase, (**Fig.3b**). In stark contrast, when a similar comparison of 2 DPI vs 4 DPI was tested for enrichment in animals receiving baricitinib, we observed negative enrichment, indicating that inflammatory genes were expressed at lower levels already after only 2 days of treatment **(Fig.3b)**. To confirm the robustness of our enrichment analysis in detecting downregulation of inflammatory pathways with treatment, we also conducted GSEA analyses using direct cross-sectional comparisons (i.e. 4 DPI untreated vs 4 DPI treated); these data demonstrated that inflammatory signatures were significantly lower in animals receiving baricitinib (4 DPI), although equivalent when comparing 2 DPI samples in which neither group had received the drug (**Fig. S5**).

To explore the impact of baricitinib on the inflammatory responses induced by SARS-CoV-2 infection at the gene level, we examined several pathways in greater detail (**Fig.3c-j**). One of the highest scoring pathways, neutrophil degranulation, was significantly enriched at 4 DPI relative to 2 DPI in the untreated group (p<0.001) (**Fig.3c**). Strikingly, however, enrichment of this pathway was completely abrogated in the treated group (p=0.979). When we examined individual genes that were (i) elevated by SARS-CoV-2 infection, and (ii) influenced by baricitinib treatment, we observed that several genes were those encoding degradative and bactericidal enzymes present in neutrophil granules (MMP9, MMP25, BPI, MPO), or highly expressed on polymorphonuclear neutrophils (CXCR1 and CXCR2), the alarmin S100A12, and genes for proteins that act to degrade the extracellular matrix during neutrophil extravasation: SERPINB10, ADAM8 (**Fig.3g**). These genes were highly upregulated in BAL samples of untreated animals, but substantially attenuated in treated animals, many at levels equivalent to pre-infection (**Fig.3g**). Collectively, these gene signatures suggest that baricitinib treatment may dampen neutrophil recruitment and activity in the lower airway during acute SARS-CoV-2 infection. We also examined the enrichment of neutrophil pathway genes in cross-sectional GSEA comparisons, as shown in **Fig. S5;** this analysis largely mirrored our D2 vs D4 observations; as many of the enriched genes were similar. Additionally, we observed several alarmin proteins (S100A8, S100A9) had lost their induction at 4 DPI in animals receiving baricitinib, as did the MPO gene. These genes have been recently demonstrated to be highly expressed in the myeloid compartment of peripheral blood of patients exhibiting severe COVID-19 disease (Schulte-Schrepping et al., 2020; Silvin et al., 2020). Baricitinib treatment also rapidly induced near complete abrogation of inflammation mediators downstream of TNFα signaling and IL6 signaling (**Fig.3 d and e, h and i**). Within these pathways, amongst the molecules suppressed by baricitinib were chemotactic factors critical for recruitment of neutrophils (CXCL6, CXCL3) and macrophages (CCL2), inflammatory serine protease factors (SERPINB2, TNFAIP6) and cytokines regulating inflammation and immune responses (IL12B). Of note, genes identified as upregulated in rheumatoid arthritis (RA) were found to be significantly enriched (p=0.0448) in untreated as compared to treated animals at 4 DPI, despite similar gene expression at 2 DPI (**Fig. S6**). In the leading-edge analysis of the rheumatoid arthritis (RA) pathway we noted lower expression of several inflammatory mediators such as CXCL8, IL1B, CCL5, CCL3, CCL20, IL18, IL6 and CXCL12 (**Fig. S6**). As baricitinib was developed to ameliorate inflammation in RA by inhibiting JAK1/2 signaling, and consistently with the reduction in the IL-6/JAK/STAT3 signaling pathway (**Fig. 3i**), these data confirm the effectiveness of baricitinib in the lower airway of SARS-CoV-2 infected RMs.

Several of the genesets comprised of genes in Type I interferon signaling and interferon stimulated genes (ISGs) presented an elevated expression relative to baseline (**Fig.3j**). In both treated and untreated groups, we observed a slight reduction in expression at 4 DPI relative to 2 DPI (**Fig 3b and f**) However, unlike genesets associated with inflammation, genes associated with Type I IFN signaling and innate antiviral responses were unperturbed by baricitinib treatment. Collectively, these data demonstrated that baricitinib treatment potently suppressed inflammatory pathways in the lower airway of RMs infected with SARS-CoV-2, but left innate antiviral signaling largely intact.

**Figure 3.**
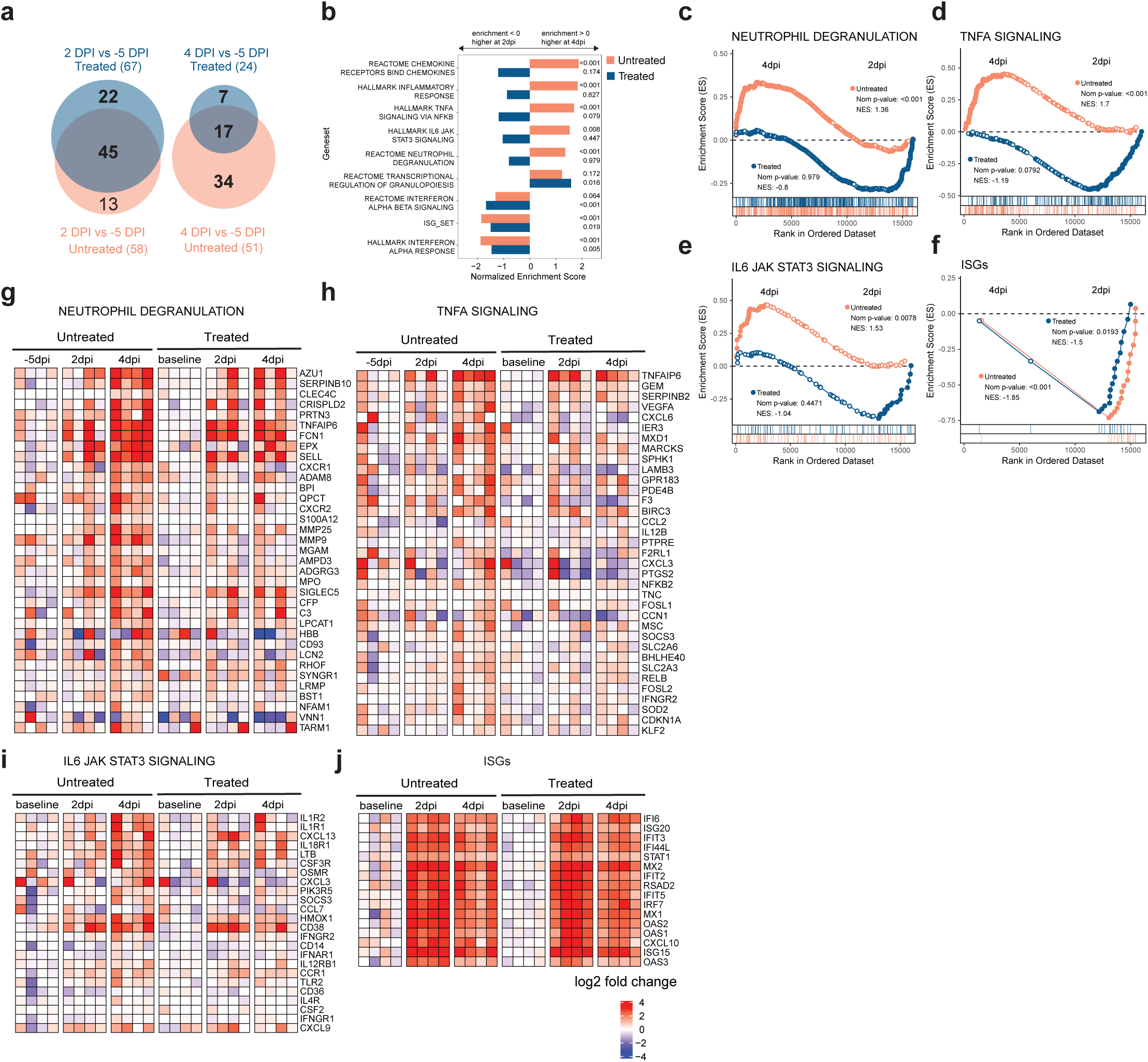
Baricitinib treatment suppresses gene expression of inflammation and neutrophil degranulation in the BAL of SARS-CoV-2 infected NHPs. Bulk RNA-Seq profiles of BAL cell suspensions from NHPs obtained at Day -5 prior to SARS-CoV-2 inoculation (Baseline), at 2 days post-inoculation, prior to baricitinib administration (2 DPI) treatment, and at 4 days post-inoculation, 2 days after initiation of baricitinib treatment (4 DPI). Both the untreated arm and baricitinib arm were comprised of n = 4 animals. (**a**) Venn diagrams indicating the number of DEGs detected at 2 DPI or 4 DPI relative to -5dpi in the untreated (blue) and baricitinib treated (red) experimental groups. The total DEGs for each comparison are shown in parentheses. (**b**) Bar plots showing enrichment of top scoring inflammatory and immunological gene signatures from the MSIGDB (Hallmark and Canonical Pathways) and databases, and custom genesets (ISGs, see below) ranked by GSEA comparisons of gene expression in the 4 DPI vs 2 DPI samples from the untreated animal group (pink bars), or 4 DPI vs 2 DPI in the baricitinib treated group (blue bars). The x-axis depicts the normalized enrichment score (NES); a positive enrichment score indicated higher expression at 4 DPI relative to 3 DPI, conversely, negative scores of a pathway indicate cumulatively higher expression in 2 DPI samples relative to 4 DPI. Nominal p-values are indicated; negative enrichment (2 DPI > 4 DPI) is indicated by bars facing left, positive enrichment (4 DPI > 2 DPI) denoted by bars facing right. (**c-f**) GSEA enrichment plots depicting pairwise comparison of gene expression of 2 DPI samples vs 4 dpi samples. 2 DPI vs 4 DPI comparisons for the untreated group are depicted by red symbols, and comparisons for the baricitinib treated group are shown in blue. The top-scoring (i.e. leading edge) genes are indicated by solid dots. The hash plot under GSEA curves indicate individual genes and their rank in the dataset. Left-leaning curves (i.e. positive enrichment scores) indicate enrichment, or higher expression, of pathways at 4 DPI, right-leaning curves (negative enrichment scores) indicate higher expression at 2 DPI. Sigmoidal curves indicate a lack of enrichment, i.e. equivalent expression between the groups being compared. The normalized enrichment scores and nominal p-values testing the significance of each comparison are indicated. (**c**) REACTOME_ NEUTROPHIL_DEGRANULATION (MSIDB #M27620) (**d**) GSEA line plot of HALLMARK_TNFA_SIGNALING_VIA_NFKB pathway (MSIGDB # M5890). (**e**) GSEA line plot of HALLMARK_IL6_JAK_STAT3_SIGNALING (MSIGDB# M5897) (**f**) a custom geneset of ISGs from prior NHP studies (Nganou-Makamdop et al., 2018; Palesch et al., 2018; Sandler et al., 2014); (**g-h**) Heat maps of top-scoring (i.e. leading edge) from the untreated 4 DPI vs 2 DPI GSEA analyses. The color scale indicates the log2 expression relative to the median of all baseline samples.

### Baricitinib treatment abolishes inflammatory cytokine and neutrophil chemoattractant expression in bronchoalveolar macrophages of SAR-CoV-2 infected rhesus macaques

The bulk RNA-Seq data indicated that gene signatures consistent with neutrophil infiltration and cytokine driven inflammation were evident as early as 2 DPI, and that baricitinib was capable of abrogation of these pathways in as quickly as 48 hours. To identify the cellular component orchestrating airway inflammation, we performed single-cell RNA-Seq (sc-RNA-Seq) profiling using 10X Genomics-based droplet sequencing. Single cell suspensions of BAL samples from three untreated and two baricitinib treated animals prior to infection, and at 4 DPI were subjected to 10X droplet capture within 3 hours of collection. After processing to remove erythrocytes and low-quality cells, the captures yielded a cumulative 45,583 cells across all samples for analysis. The cellular distribution is summarized in the UMAP shown in **Fig.4a**. Similar to observations reported in sc-RNA-Seq data in humans infected with SARS-CoV-2 (Bost et al., 2020; Chua et al., 2020; Liao et al., 2020), the vast majority of cells in BALs were predominantly macrophage/myeloid origin (80.7%), followed by lymphocytes (CD4^+^/CD8^+^ T cells/ NK cells) (9.8%) and approximately 3.2% were identified as epithelial. Allocation of cells from the cumulative data by treatment variables (**Fig.4b**) demonstrated that the cellular distribution was distributed equivalently amongst the experimental groups and no population was enriched due to batch or technical variation associated with individual captures. We probed the macrophage population for upstream regulators associated with the inflammatory pathways identified in the bulk RNA-Seq analyses, and observed elevated expression of several inflammatory mediators at 4 DPI: IL6, TNF*α*, IL1*β* and IL10 (**Fig.4c; Fig. S7 and S8)**. IFN*β* was also highly expressed in the macrophage cluster, however, IFN*α* transcripts were detected in a virtually negligible fraction of cells **Fig.4c; Fig. S7 and S8; and data not shown**). Strikingly, and consistent with the bulk RNA-Seq data, we observed that baricitinib treatment virtually abrogated expression of TNF*α*, IL10, IFN*β* and IL6 in pulmonary macrophages, and significantly reduced expression of IL1*β* (**Fig.4c**). We also observed a robust induction of chemokines driving neutrophil recruitment (CXCL3/MIP2*β*, CXCL8/IL8), macrophage trafficking (CCL4L1/MIP1*β*), and CXCL10/IP10 (**Fig.4d and e**), a pleiotropic chemokine upregulated in several viral infections, and long hypothesized to be associated with pathogenesis in SARS-CoV viral infection and observed in SARS-CoV-1 infection of NHPs(de Lang et al., 2007; Laing et al., 2020; Tang et al., 2005). Notably, after 48 hours of baricitinib treatment, expression of these proinflammatory cytokines was reduced to negligible levels (**Fig.4d and e**). Examination of the expression levels of antiviral ISGs in pulmonary macrophages yielded a much different pattern than those observed for inflammatory genes – although widespread induction of ISGs were observed after SARS-CoV-2 infection, baricitinib treatment had only a very modest impact on these pathways (**Fig.4f**). Collectively, these data support a model in which baricitinib administration strongly reduces airway inflammation and neutrophil accumulation, but has a minimal effect on innate antiviral immunity.

**Figure 4.**
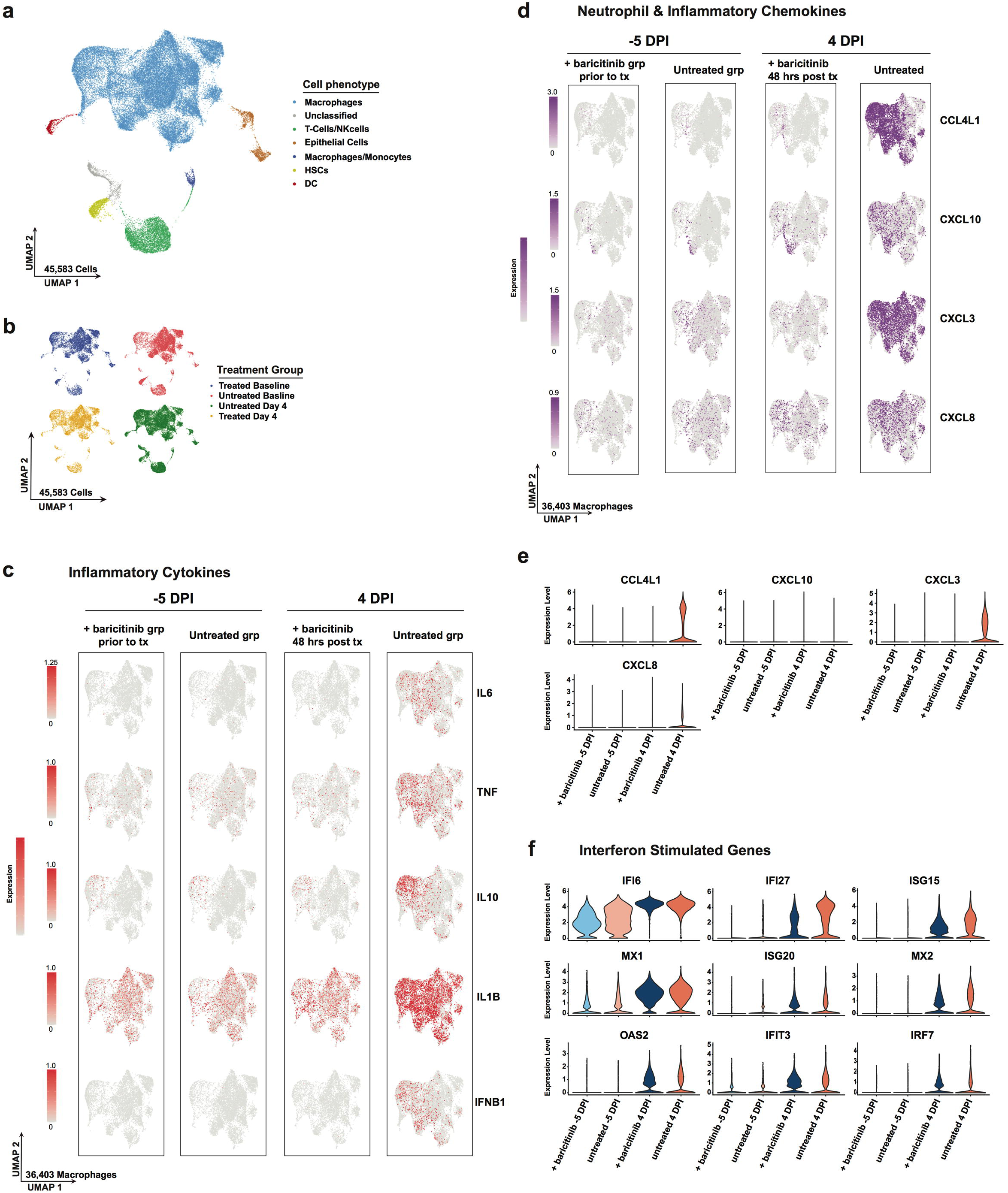
Baricitinib treatment abolishes inflammatory cytokine and neutrophil chemoattractant expression in bronchoalveolar macrophages. Single cell suspensions from BALs of SARS-CoV-2 infected RMs were subject to 10X Genomics capture and sequencing. (**a**) UMAP showing major cell types in BAL samples (n = 10 samples; untreated, baseline n = 3; untreated, 4 DPI n = 3; treated, baseline n = 2; treated, 4 DPI). (**b**) UMAP showing clusters in BAL samples by treatment days (n = 10). (**c**) UMAP projection of pro-inflammatory cytokines in macrophages. (**d**) UMAP projection of neutrophil chemoatttractants and pro-inflammatory chemokines. (**e** and **f**) Expression of chemokines and interferon stimulated genes (ISGs) in treated and untreated samples at baseline and 4 DPI. The colored expression scale of expression in UMAPs is depicted on a per gene basis: the scale represent the per cell reads for each gene divided by the total reads for of that cell, scaled to the factor shown and natural log-transformed.

### Baricitinib leads to reduced BAL levels of neutrophils and neutrophil NETosis activity

To gain insight into the immunologic effects of baricitinib treatment on cellular distribution within BAL, we applied global high-dimensional mapping of 23-parameter flow cytometry data. As shown in the UMAP representation (**Fig.5a**), untreated and baricitinib treated RMs had different BAL cellular distribution starting from 4 DPI, including in neutrophils. This was of particular interest considering the higher expression of neutrophil-attracting chemokines in untreated RMs (**Fig.4d and e**), thus we focused our flow cytometry immunologic analyses in determining longitudinal levels of neutrophils (CD45^+^CD3^-^CD20^-^CD66^+^ live granulocytes; representative staining in **Fig. S9a**, and **Fig.5b**). Analyses of BAL showed a rapid recruitment of neutrophils in the lung at 4 DPI during the peak of viremia. All untreated animals showed sustained, higher frequencies of neutrophils at later stages of infection (10-11 DPI) as compared to baricitinib treated animals (**Fig.5c and d;** p=0.0286). In blood, neutrophils (**Fig.5e**) remained relatively stable post infection as compared to pre-infection and at lower levels in untreated as compared to treated animals at the latest experimental points (p=0.0571), consistently with a higher migration to lung in untreated RMs. Since the flow cytometry data of BAL shows a reduced migration of neutrophils to lung in baricitinib-treated animals, we next measured neutrophil extracellular trap (NET) activity by quantification of extracellular DNA via Sytox staining, a functional readout of NETosis activity (**Fig.5f and g**). NETs have been reported as an important mechanism of inflammation and microvascular thrombosis in patients with COVID-19 (Skendros et al., 2020). Baricitinib treated RMs showed decreased NET formation by blood neutrophils at 4 and particularly 10 DPI when compared to untreated animals (**Fig.5g;** p=0.0571).

**Figure 5.**
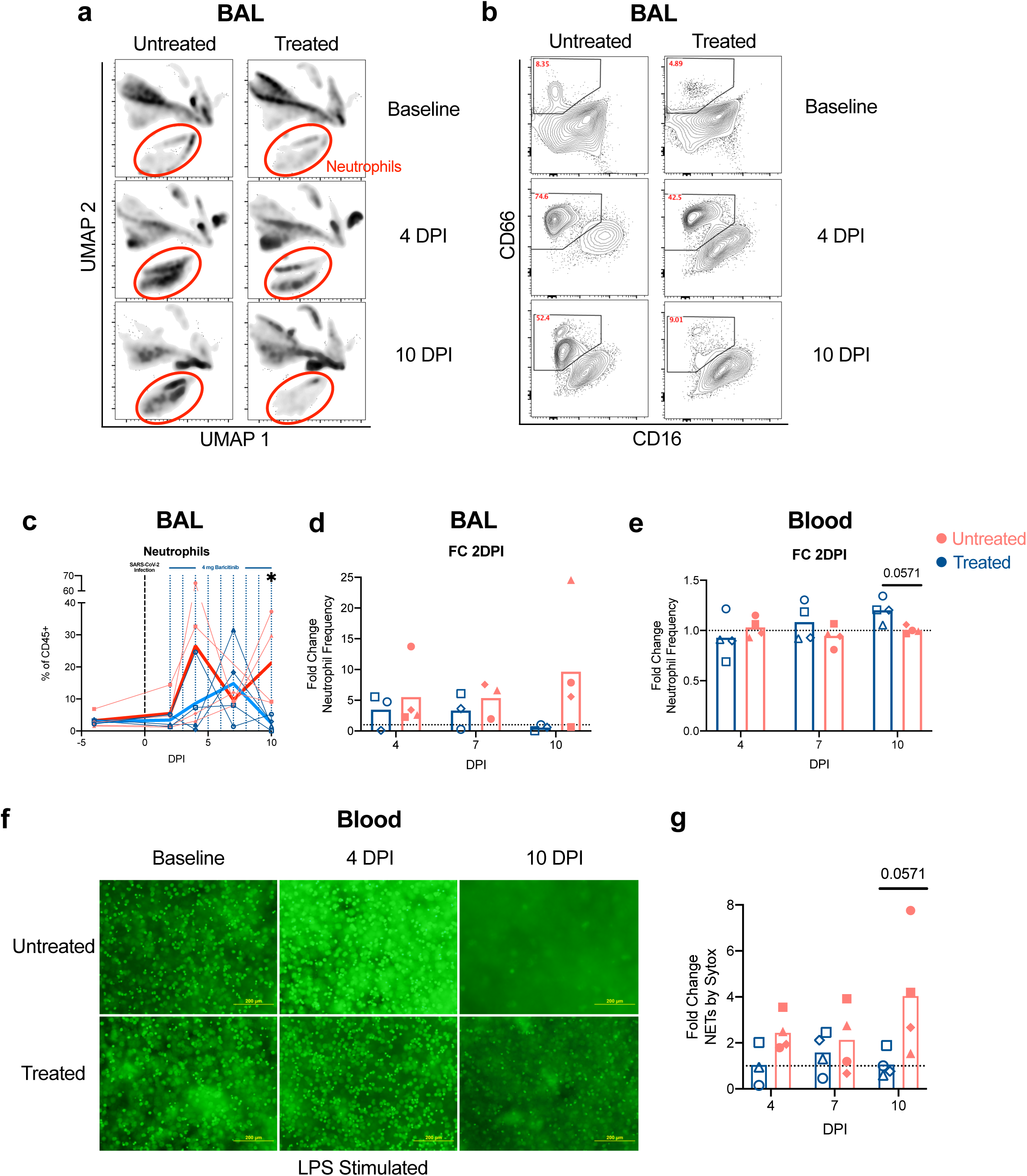
Baricitinib treated RMs have decreased infiltration of innate immune cells and lowered neutrophil NETosis. (**a**) UMAP analysis of BAL in baricitinib treated (n= 4) and untreated (n= 4) SARS-CoV-2 infected RMs before infection (D -5 PI; baseline), and at 4, and 10 DPI. (**b**) Representative flow cytometry staining of neutrophil infiltration in BAL at baseline, and 4 and 10 DPI. Longitudinal levels of neutrophils within BAL samples depicted as a % of CD45+ cells and fold change to 2 DPI (**d**) in treated (blue) and untreated (red) RMs. (**e**) Fold change to 2 DPI of neutrophils in blood of baricitinib treated and untreated SARS-CoV-2 infected RMs. (**f**) Representative microscopy images of NETS by Sytox green assay in baricitinib treated and untreated SARS-CoV-2 infected RMs. Scale bar: 200 μm. (**g**) Quantification of NETosis activity upon staining extracellular DNA with Sytox in isolated stimulated neutrophils from blood. Fold change of Sytox levels to -5 DPI. In c, d, e, and g each symbol represents individual animals. Thick lines represent the average of the baricitinib treated (blue line), and untreated groups (red line). Bars in d, e, and g represent the average of the treated and untreated groups. Statistical analysis was performed using a non-parametric Mann-Whitney Test.

Altogether, these data support baricitinib activity in reducing inflammation by decreasing pro-inflammatory neutrophilic levels, activity and NETosis.

### Baricitinib reduced T cell immune activation in SARS-CoV-2 infected rhesus macaques

Our transcriptomic data indicated that baricitinib reduced the expression of multiple cytokines that can induce T cell immune activation. As such, we then analyzed levels of T cells, and their frequency of activation and proliferation by flow cytometry (gating strategy shown in **Fig. S9b**). CD4^+^ T cell levels in blood remained similar between treated and untreated animals, with 1/4 baricitinib treated and 2/4 untreated RMs exhibiting a pronounced reduction in CD4^+^ T cell frequencies at 10 DPI (**Fig.6a**). When analyzing different CD4^+^ T cell subsets, we observed an expansion of CD4^+^ T_Regs_ at 4 (p=0.0571) and 6 DPI in the untreated, but not in the baricitinib treated RMs, both as frequency (**Fig.6b and c**) and absolute numbers (not shown). Specifically, the mean fold change in CD4^+^ T_Regs_ frequency at 4 and 6 DPI, as compared to pre-treatment baseline (2 DPI), was of 7.43 and 4.36 in untreated and of 1.22 and 1.13 in baricitinib treated RMs, respectively, suggesting higher levels of inflammation in the untreated group resulting in greater expansion of CD4^+^ T_Regs_. Peripheral CD8^+^ T cells were reduced at 10 DPI in 2/4 baricitinib treated and 2/4 untreated animals (**Fig.6d**). Differences in immune activation and proliferation were evident among treated and untreated animals for CD8^+^ T cells as well. Specifically, the frequency of proliferating (Ki-67^+^) memory CD8^+^ T cells in blood progressively and significantly increased in all 4 untreated animals at 7 and 10 DPI, while significantly decreasing in all baricitinib treated RMs already at 4 DPI. As a result, at 10 DPI the mean frequency of CD8^+^Ki-67^+^ was significantly higher in untreated animals (24.38% vs 7.38%; p = 0.0286, **Fig.6e**).

**Figure 6.**
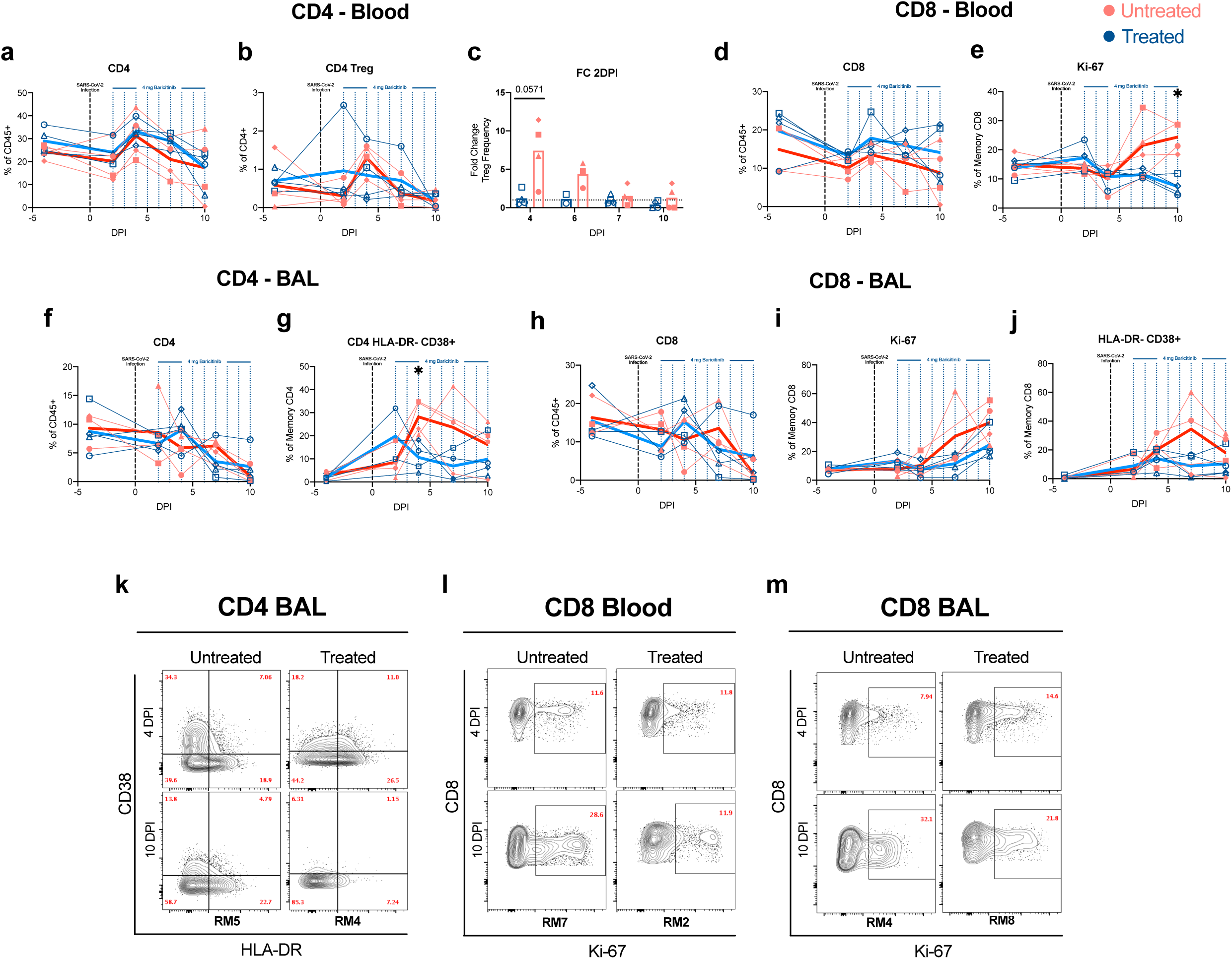
Decreased levels of T cell proliferation and activation in baricitinib treated RMs. Longitudinal levels of (**a**) circulating CD4^+^ T cells and (**b**) CD4^+^ T_Reg_ cells measured by flow cytometry of baricitinib treated (blue) and untreated (red) SARS-CoV-2 infected RMs. (**c**) Fold changes to 2 DPI of circulating CD4^+^ T_Reg_ cells. (**d**) Levels of circulating CD8^+^ T cells and (**e**) proliferating (Ki-67^+^) memory CD8^+^ T cells. (**f**) Levels of CD4^+^ T cells, and (**g**) HLA-DR^-^CD38^+^ CD4^+^ T cells in bronchoalveolar lavages (BAL) measured by flow cytometry. (**h**) Levels of CD8^+^ T cells, (**i**) proliferating (Ki-67^+^) memory CD8^+^ T cells and (**j**) HLA-DR^-^CD38^+^ CD8^+^ T cells in BAL. Each symbol represents individual animals. Thick lines represent the average of the baricitinib treated (blue line), and untreated groups (red line). (**k-m**) Representative staining of Ki-67 and CD38 by HLA-DR. Bars in (c) represent the average of the treated and untreated groups. Statistical analysis in b was performed using non-parametric Mann-Whitney Test.

CD4^+^ T cells in the BAL remained relatively constant until 7 DPI, when the majority of animals started experiencing a reduction in their frequencies (**Fig.6f**). CD4^+^ T_Regs_ expanded more at 4 and 7 DPI in the untreated group, again hinting at higher levels of inflammation and trafficking of expanded T_Regs_ from blood to lung (data not shown). Consistently, untreated RMs showed an early (present at 4 DPI), large (mean fold change of 3.31 at 7 DPI vs 2 DPI compared to 1.14 in the treated RMs) and prolonged (up to 10 DPI) increase in the frequency of memory CD4^+^ T cells expressing CD38 (CD38^+^HLA-DR^-^; 4 DPI, p=0.0286, **Fig.6g**); similar results were found for memory CD4^+^ T cells expressing CD38 and HLA-DR (CD38^+^HLA-DR^+^; not shown). Remarkably, different from untreated RMs, the frequency of those activated memory CD4^+^ T cells decreased in baricitinib treated animals starting at 4 DPI and remained lower than pre-treatment until 10 DPI (**Fig.6g)**. Consistent with a reduced pro-inflammatory state of CD4^+^ T cells, baricitinib treated RMs showed a lower frequency of CD4^+^ T cells that spontaneously (without stimulation) produced pro-inflammatory, Th17 related cytokines (IL-17^+^; IL-17^+^IL-21^+^; IL-17^+^IL-22^+^) when compared to untreated RMs (**Fig. S10**).

As with CD4^+^ T cells, the reduction in CD8^+^ T cells was more pronounced in BAL, starting at 7 DPI and maintained until necropsy (**Fig.6h**). Similarly, also in BAL the frequency of CD8^+^Ki-67^+^T cells increased more extensively in untreated than baricitinib-treated RMs (30.53% vs 11.53% at 7 DPI; 39.95% vs 24.65% at 10 DPI; **Fig.6i**); as a result, the fold change (as compared to 2 DPI, pre-treatment) in the frequency of memory CD8^+^Ki67^+^ T cells was higher in untreated then baricitinib treated animals both at 7 (8.22 vs 1.02) and 10 (6.28 vs 2.48) DPI. A similar trend was measured for activated memory CD8^+^ T cells, with higher frequency and fold change in untreated than baricitinib treated animals at 7 (FC CD38^+^DR^-^: 23.67 vs 1.62; CD38^+^DR^+:^ 4.55 vs 0.90) and 10 (FC CD38^+^DR^-^: 9.81 vs 1.43; CD38^+^DR^+^: 16.66 vs 2.90) DPI (**Fig.6j**, and data not shown). Representative staining for Ki-67 and HLA-DR by CD38 in CD4^+^ and CD8^+^ T cells are shown in Fig.6k-m.

Finally, we assessed the ability of peripheral T cells to respond to *ex vivo* SARS-CoV-2 specific stimulation (with a SARS-CoV-2 S peptide pool characterized in (Grifoni et al., 2020)) and to non-antigen specific stimulation (with PMA/ionomycin). Importantly, the levels of SARS-CoV-2 specific CD4^+^ and CD8^+^ T cells producing IFNγ, TNFα, IL-2, IL-4 and IL-17a in response to S peptide pool stimulation were similar in both groups of animals (**Fig. S11a-c**). Similarly, the frequency of CD4^+^ and CD8^+^ T cells producing IL-17a, IL-21, IL-22, IFNγ, and TNFα were similar among the two groups after PMA/Ionomycin stimulation (**Fig. S11 d and e**). Furthermore, levels of memory CD4^+^ and CD8^+^ T cells expressing granzyme B or PD-1 remained very similar between untreated and treated RMs both in blood (**Fig. S11 f and g**) and BAL (**Fig. S11 h and i**).

Collectively, these findings indicate that baricitinib treatment lead to downstream reduction in T cell activation and proliferation, without an overall detrimental effect to antiviral function of T cells.

## Discussion

In this study, we tested baricitinib, a JAK1/2 inhibitor clinically approved for rheumatoid arthritis, as a therapeutic candidate to reduce systemic inflammation caused by SARS-CoV-2 infection in RMs. Notably, baricitinib treated RMs displayed reduced (i) lung pathology, from moderate in untreated animals to mild; (ii) levels of inflammatory cytokines, chemokines, and signaling pathways associated with inflammation, neutrophil recruitment, and disease progression in SARS-CoV-2 infected humans; and (iii) levels of systemic inflammation that are associated with COVID-19 severity in humans while not having an impact on Type 1 IFN responses. This beneficial anti-inflammatory effect of baricitinib was confirmed by a reduced infiltration of neutrophils into the lungs, and a highly reduced T cell activation in both blood and BAL as compared to untreated animals. Furthermore, in this study we were able to observe an increased NETosis activity of neutrophils upon SARS-CoV-2 infection, previously described in serum from COVID19 patients (Skendros et al., 2020), which was reduced in baricitinib treated animals. Remarkably, single-cell RNA sequencing showed reduced immune activation, neutrophil recruitment, and macrophage trafficking signatures in pulmonary macrophages from treated RMs already after two doses of baricitinib, at 4 DPI. IL-6, TNFα, IL-10, IL-1B, CXCL3/MIP-2*β*, CXCL8/IL8, CCL4L1/MIP-1*β*, and CXCL10/IP-10 were all expressed at higher levels in pulmonary macrophages from untreated animals compared to baricitinib treated animals. These data confirm very recent studies that demonstrated by RNA-Seq analysis that higher levels of inflammatory cytokines in lung macrophages are associated with patients presenting with severe/critical COVID-19 cases (Liao et al., 2020). Thus, baricitinib could have clinical benefits in reducing the inflammatory response typically seen in moderate to severe cases of COVID-19 (**Fig. 7**). Of note, one of the advantages of baricitinib when compared with other cytokine-specific anti-inflammatory therapies is that it can be more effective by inhibiting several cytokines involved in the cytokine storm described in severe cases of COVID-19.

**Figure 7.**
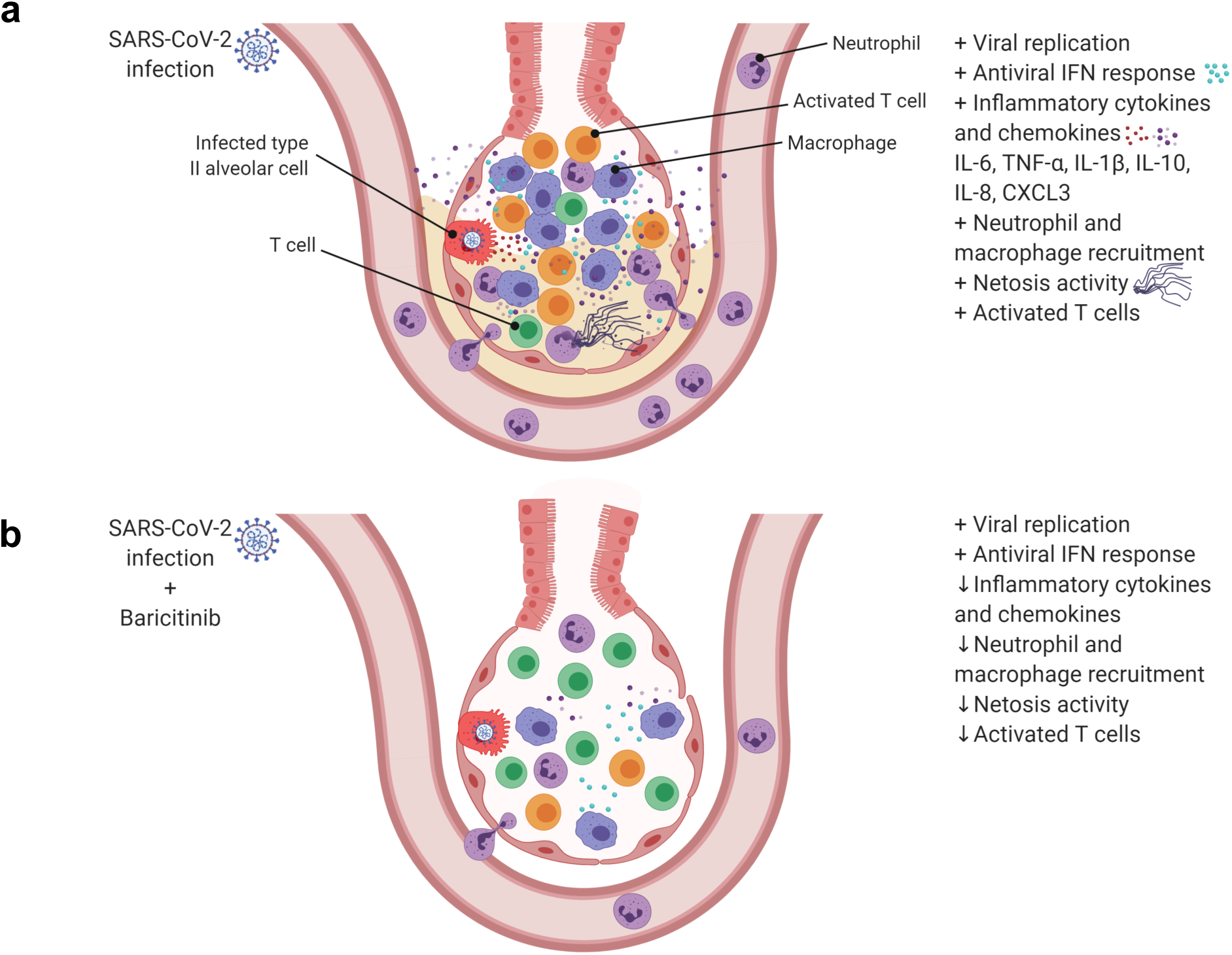
(a) SARS-Cov-2 infection in rhesus macaques results in an accumulation of inflammatory macrophages and neutrophils in the lower airway. These airway macrophages produce high amounts of inflammatory cytokines and neutrophil-attracting chemokines and show upregulated Type I interferon signaling. Neutrophil NETs and the inflammation induced by SARS-CoV-2 infection both contribute to lung pathology. (**b**) Baricitinib treatment reduced the levels of macrophages producing inflammatory cytokines and neutrophil-attracting chemokines, decreased the infiltration of neutrophils into the lung and reduced T cell activation. The Netosis activity of neutrophils was also reduced. In treated animals, the antiviral interferon response was maintained, viral replication was not impacted, and lung pathology was mild.

Clinical pathology and laboratory parameters of toxicity remained similar in the treated animals for the 8-9-day treatment course at a dose equivalent to humans (4 mg daily; (Bronte et al., 2020; Cantini et al., 2020; Titanji et al., 2020)). Baricitinib was found distributed in lungs, a key tissue for SARS-CoV-2 replication, as well as in the central nervous system (CNS). Although several *in silico* modeling and *in vitro* studies suggested baricitinib as a possible treatment candidate to COVID-19 due to its potential antiviral activity (Cantini et al., 2020; Richardson et al., 2020; Stebbing et al., 2020; Titanji et al., 2020), we did not observe changes in viral replication dynamics in the treated animals. One of the main concerns in using a JAK inhibitor such as baricitinib, is that its downstream anti-immune activation effects could limit immune responses necessary to combat SARS-CoV-2. Importantly, we did not identify reduction of SARS-CoV-2 specific and unspecific CD4^+^ and CD8^+^ T cell responses in treated animals, indicating that baricitinib reduces systemic inflammation without any detrimental effects to the adaptive antiviral immune responses. Baricitinib did not inhibit genes associated with Type I Interferon antiviral responses, indicating its mode of action in this context is primarily to dampen inflammatory responses and neutrophil recruitment while maintaining innate and adaptive antiviral pathways.

Several ongoing clinical trials are studying the effects of baricitinib in SARS-CoV-2 infected humans. Currently, the Adaptive COVID-19 Treatment Trial (ACTT-2; NCT04401579) is evaluating the effects of baricitinib in combination with remdesivir and the COV-BARRIER trial (NCT04421027) is studying baricitinib as a monotherapy. Participants are treated with placebo, remdesivir alone or a combination of remdesivir and baricitinib. In a retrospective cohort study, 15 patients with severe COVID-19 were administered a short-course of baricitinib in combination with hydroxychloroquine and this was associated with a complete recovery in 11/15 subjects(Titanji et al., 2020). In a separate pilot study, baricitinib was combined with lopinavir-ritonavir in 12 patients starting treatment 6 days post-symptom onset, with all individuals showing significantly improved clinical and laboratory parameters with no treated individuals requiring ICU care (Cantini et al., 2020). Being performed in an animal model, this study has some key advantages and some important limitations. Advantages include the ability to correct for parameters that may impact clinical outcome and treatment readout, including using the same virus inoculum, dose, and route of infection, and starting baricitinib at the same phase of infection in all subjects. Furthermore, the NHP model permits longitudinal collection of BAL and lung at necropsy, which in turn allows the in-depth characterization of the impact of baricitinib on immune activation and immunologic responses, including single-cell RNA sequencing analyses of macrophages, at the foci of infection. The main limitation of the study is the small group size, with a total of eight RMs; furthermore, being that SARS-CoV-2 infected RMs is a model of moderate COVID-19 (at least in animals between 11-17 years old as in our study) and with treatment initiated early after infection, we cannot determine if the therapeutic impact of baricitinib will be the same in severe COVID-19 patients and when started later than 2 days post-infection. Mitigating that concern, in our study, treatment started at peak viremia, including in BAL, and once inflammatory signatures were already highly up-regulated, mimicking the conditions in which baricitinib would be administered clinically. Indeed, a recent small clinical trial in 20 severe COVID-19 patients indicated that baricitinib mitigated immune dysregulation by reducing plasma levels of IL6, IL1*β* and TNF*α* and lowered time to recovery(Bronte et al., 2020). Our data provides rationale for baricitinib treatment in COVID-19 to be given in a window where blocking immune inflammation would prevent the formation of a cytokine storm without interfering in the initial responses necessary for preventing viral dissemination and persistence.

In conclusion, this study provides rationale and mechanisms of actions for a beneficial anti-inflammatory effect of baricitinib treatment for COVID-19.

## Acknowledgments

We kindly thank the Yerkes National Primate Research Center (YNPRC) Division of Animal Resources, especially Stephanie Ehnert, Stacey Weissman, Denise Bonenberger, John M. Wambua, Dominic M. D’Urso, Racquel Sampson-Harley and Kalpana Patel in Research Resources and Joyce Cohen in Veterinary Medicine, for providing support in animal care. Pharmaceutical-grade Baricitinib was commercially obtained, provided free of charge by Dr. Schinazi, and it was found to be > 99% pure by LC-MS-MS. This study was primarily supported by an Emory University COVID-19 Molecules and Pathogens to Populations and Pandemics (MP3) Initiative Seed Grant to M.Pa., A.P., and R.F.S., and by YNPRC Coronavirus Pilot Research Project Program grant to M.Pa. under award P51 OD11132. This work was supported by the NIAID, NIH, under awards R37AI141258, R01AI116379 to M.Pa. and R01MH116695 to R.F.S and U24 AI120134 to S.E.B. Support for this work was also provided by award NIH Office of Research Infrastructure Programs (ORIP) P51 OD11132 to YNPRC, P51OD011092 to ONPRC, 1S10OD025002-01 to the Integrated Pathology Core/ONPRC, and NIH National Institute of Allergy and Infectious Diseases (NIAID) award P30 AI050409 to the Center for AIDS Research (CFAR) at Emory University. Next generation sequencing services were provided by the Yerkes NHP Genomics Core which is supported in part by NIH P51 OD011132. Sequencing data was acquired on an Illumina NovaSeq6000 funded by NIH S10 OD026799 to S.E.B. This work was additionally funded by grants from the National Institutes of Health/National Institutes of Allergy and Infectious Diseases R01 AI143411-01A1 and R01 AI149672 to J.D.E. The content of this publication does not necessarily reflect the views or policies of the U.S. Department of Health and Human Services, nor does it imply endorsement of organizations or commercial products.

## Author Contributions

Conceptualization, T.H., M.Pi., J.L.H., A.P., S.E.B., R.F.S., and M.Pa.; Methodology, T.H., M.Pi., A.K.B., E.G.V., K.B., A.A.U., G.K.T., K.L.P., S.G., S.K., S.T., O.D., K.A.C., M.N.S., L.W., P.D.F., J.W., A.P., S.P.K., C.E.S., S.W., H.A., E.A.M., M.YH.L., K.Z., S.T., T.R.H., E.N.B., S.P.R., T.H.V.; Formal Analysis, T.H., M.Pi., Z.S., E.G.V., A.K.B., K.B., A.A.U., G.K.T., S.G., S.K., S.T., P.D.F., J.W., A.P., S.P.K., S.P.R., T.H.V; Investigation, T.H., M.Pi., E.G.V., J.C., S.J., J.S.W., F.C-S., R.L.S., R.D.L., A.P., S.P.R., R.P.S., and T.V.; Resources, D.W., R.F.S., S.B., and M.Pa.; Writing – Original Draft, T.H., M.Pi., J.L.H. and M.Pa.; Writing – Review & Editing, T.H., M.Pi., J.L.H., S.B. and M.Pa.; Visualization, T.H., M.Pi., A.K.B., A.A.U., G.K.T., Z.S. and E.V.; Supervision, R.F.S., S.E.B., and M.Pa.; Funding Acquisition, T.V., A.P.,S.E.B. R.F.S., and M.Pa.

## Competing Interests Statement

Dr. Raymond Schinazi served as an unpaid consultant for Eli Lilly whose drugs are being evaluated in the research described in this paper. In addition, Dr. Schinazi owns shares in Eli Lilly. The terms of this arrangement have been reviewed and approved by Emory University in accordance with its conflict of interest policies. All other authors do not have any conflicts to declare.

## Methods

### Viral Stocks

Vero E6 cell line (African Green Monkey Kidney cell line; CRL-1586, ATCC) was used in this study. Vero cells were cultured and maintained in MEM (Sigma) supplemented with 10% heat inactivated fetal bovine serum (FBS) (Gibco) and 1 mM L-glutamine (Gibco), 50 U/ml penicillin and 50 μg/ml streptomycin (Gibco). The cells were kept at 37°C in the presence 5% CO2. At the time of virus inoculation and propagation, the concentration of FBS was reduced to 2%. SARS-CoV-2 (NR-52281: BEI Resources, Manassas, VA; USA-WA/2020, Lot no. 70033175) was passaged on Vero E6 cells at a MOI of 0.01 to produce the infectious viral stock. SARS-CoV-2 has been propagated and titrated by TCID_50_ method followed by storage of aliquots at -80°C until further use in the experiments.

Back titration of viral stocks via plaque assay was used to determine the infectious dose delivered to the RMs. The virus stock was also directly sequenced via metagenomic methods prior to inoculation to confirm the presence of the furin cleavage motif, which has been shown to be lost upon sequential passage of SARS-CoV-2 in culture (Davidson et al., 2020). Our stock contained fewer than 6% of viral genomes with a mutation that could potentially abrogate furin-mediated cleavage of S (data not shown).

### Study Design

Eight specific-pathogen-free (SPF) Indian-origin rhesus macaques (RM; *Macaca mulatta;* **Supplementary table 1**) were housed at Yerkes National Primate Research Center (YNPRC) as previously described (McGary et al., 2017). Animals for study assignment were requested to be greater than 11 years old without preference for gender or MHC haplotype. RMs were infected with 1.1x10^6^ plaque forming units (PFU) SARS-CoV-2 via both the intranasal (1 mL) and intratracheal (1 mL) routes concurrently. Absent further stratification criteria, four RMs were administered 4 mg Baricitinib (Olumiant*®*, Eli Lilly) starting at day 2 post-infection (DPI) for 8-9 consecutive days. Baricitinib was supplied as a powder that was folded into food items (i.e. honey, yogurt, etc.) or distilled water, which was delivered either orally or as a gavage when animals were being anesthetically accessed, respectively. At each anesthetic access pulse oximetry was recorded and RMs were clinically scored for responsiveness and recumbency; discharges; skin condition; respiration, dyspnea, and cough; food consumption; and fecal consistency (**Supplementary Tables 2 and 3**). At 10-11 DPI, RMs were administered Baricitinib and subjected to necropsy after 2 hours with blood and cerebrospinal fluid (CSF) collected perimortem to assess pharmacokinetics of baricitinib. Longitudinal tissue collections of peripheral blood (PB); axillary or inguinal lymph node (LN) biopsies; bronchoalveolar lavage (BAL); and nasal, and pharyngeal mucosal swabs in addition to thoracic X-rays (ventrodorsal and right lateral views) were performed immediately prior to Baricitinib administration as annotated (**Fig.1a**). In addition to the tissues listed above, at necropsy the following tissues were processed for mononuclear cells: hilar LN, lower lung, and upper lung. Additional necropsy tissues harvested for histology included nasopharynx.

### Study Approval

YNPRC’s animal care facilities are accredited by both the U.S. Department of Agriculture (USDA) and by the Association for Assessment and Accreditation of Laboratory Animal Care (AAALAC). All animal procedures were performed in line with institutional regulations and guidelines set forth by the NIH’s Guide for the Care and Use of Laboratory Animals, 8^th^ edition, and were conducted under anesthesia with appropriate follow-up pain management to minimize animal suffering. All animal experimentation was reviewed and approved by Emory University’s Institutional Animal Care and Use Committee (IACUC) under permit PROTO202000035.

### Determination of viral load RNA

SARS-CoV-2 genomic RNA was quantified in nasopharyngeal (NP) swabs, throat swabs, plasma, and bronchoalveolar lavages (BAL). Swabs were placed in 1mL of Viral Transport Medium (VTM-1L, Labscoop, LLC). Viral RNA was extracted from NP swabs, throat swabs, and BAL on fresh specimens, while plasma was frozen for future analysis. Viral RNA was extracted manually using the QiaAmp Viral RNA mini kit according to the manufacturer’s protocol. Quantitative PCR (qPCR) was performed on viral RNA samples using the N2 primer and probe set designed by the CDC for their diagnostic algorithm: CoV2-N2-F: 5’-TTACAAACATTGGCCGCAAA-3’, CoV2-N2-R: 5’-GCGCGACATTCCGAAGAA-3’, and CoV2-N2-Pr: 5’-FAM-ACAATTTGCCCCCAGCGCTTCAG-BHQ-3’. qPCR reactions were performed in duplicate with the Thermo-Fisher 1-Step Fast virus mastermix using the manufacturer’s cycling conditions, 200nM of each primer, and 125nM of the probe. The limit of detection in this assay was 257 copies per mL of VTM/plasma/BAL. To verify sample quality the CDC RNase P p30 subunit qPCR was modified to account for rhesus macaque specific polymorphisms. The primer and probe sequences are RM-RPP30-F 5’-AGACTTGGACGTGCGAGCG-3’, RM-RPP30-R 5’-GAGCCGCTGTCTCCACAAGT-3’, and RPP30-Pr 5’-FAM-TTCTGACCTGAAGGCTCTGCGCG-BHQ1-3’. A single well from each extraction was run as above to verify RNA integrity and sample quality via detectable and consistent cycle threshold values.

### SARS-CoV-2 quantification from necropsy samples

An approximately 0.5 cm^3^ sample of each tissue was collected at necropsy, placed in 500µL Nuclisens lysis buffer (Biomerieux), and stored at -80℃. Thawed samples were homogenized with a sterile pestle, treated with 50µL proteinase K (Qiagen) for 30 minutes at 55℃, and pelleted. Total nucleic acid was extracted from 250µL of supernatant using eMAG (Biomerieux) and eluted into 50µL. RT-PCR for SARS-CoV-2 N2 was performed as previously described, and singleplex RT-PCR for RNase P was performed using primers and probes optimized for quantitation, each using 5µL of eluate (Waggoner et al., 2020). To allow for comparison of SARS-CoV-2 levels between samples that may have had subtle differences in starting material, the SARS-CoV-2 N2 Ct was normalized to the RNase P control by: 1) calculating the difference between N2 Ct and RNase P Ct for each sample, and 2) adding this to the median RNase P Ct value for the sample type. For the purposes of data visualization, samples in which SARS-CoV-2 N2 was undetected were assigned a Ct value of 40 (the assay limit of detection).

### Quantification of baricitinib by LC-MS/MS in plasma, CSF and tissue

One hundred μL of plasma or CSF samples were extracted with 500 μL of methanol. For tissues like brain and lung, 0.2 to 0.5 g of tissue were homogenized and extracted with 2 mL of methanol. [^2^H_9_]-ruxolitinib dissolved in 50% methanol at 500 nM was spiked in plasma/CSF (10 μL) or tissue samples (40 μL) as internal standard before extraction. The supernatant of each extraction (50 μL) was mixed with equal amount of 0.1% formic acid and then subjected to LC-MS/MS analysis after filtration through 0.22 μm membrane with Costar Spin-X centrifuge tube filters (Corning, NY). A Vanquish Flex HPLC system (Thermo Scientific, Waltham, MA) coupled with a TSQ Quantiva triple quadrupole mass spectrometer (Thermo Scientific, Waltham, MA) with an ESI interface was used for LC-MS analysis. Analytes were separated by a Kinetex EVO-C18 column (100 x 2.1 mm, 2.6 μm; Phenomenex, Torrance, CA) at a flow rate of 300 μL/min, 35°C. Gradient elution was used for the separation with mobile phase A (0.1% formic acid) and mobile phase B (acetonitrile). The LC gradient started with 10% of mobile phase B for 0.5 min, then increased from 10% to 90% in 4 min and kept at 90% for 0.5 min before returning to the initial condition. Selected reaction monitoring in positive mode (spray voltage: 3,200 V; sheath gas: 40 Arb; auxiliary gas: 20 Arb; ion transfer tube temperature: 350°C; vaporizer temperature: 350°C) was used to detect baricitinib (372.1 *→* 251.1) and the internal standard [^2^H_9_]-ruxolitinib (316.2 *→* 186.1). Data were collected and processed by Thermo Xcalibur 3.0 software. Calibration curves were generated from standard baricitinib by serial dilutions in blank biometric samples using the same extraction method described above. For CSF, 0.5% plasma was used as surrogate to make calibration curve. The calibration curves had r^2^ value greater than 0.99.

All the chemicals are analytical grade or higher and were obtained commercially from Sigma-Aldrich (St. Louis, MO). [^2^H_9_]-ruxolitinib was purchased from ALSACHIM (lllkirch, Alsace, France) with purity greater than 98%.

### Histopathology and immunohistochemistry

Due to study end point, the animals were euthanized, and a complete necropsy was performed. For histopathologic examination, various tissue samples including lung, nasal turbinates, trachea, or brain, were fixed in 4% neutral-buffered paraformaldehyde for 24h at room temperature, routinely processed, paraffin-embedded, sectioned at 4μm, and stained with hematoxylin and eosin (H& E). The H&E slides from all tissues were examined by two board certified veterinary pathologists. For each animal, all the lung lobes were used for analysis and affected microscopic fields were scored semi-quantitatively as Grade 0 (None); Grade 1 (Mild); Grade 2 (Moderate) and Grade 3 (Severe). Scoring was performed based on these criteria: number of lung lobes affected, type 2 pneumocyte hyperplasia, alveolar septal thickening, fibrosis, perivascular cuffing, peribronchiolar hyperplasia, inflammatory infiltrates, hyaline membrane formation. An average lung lobe score was calculated by combining scores from each criterion. Digital images of H&E stained slides were captured at 40× and 200× magnification with an Olympus BX43 microscope equipped with a digital camera (DP27, Olympus) using Cellsens® Standard 2.3 digital imaging software (Olympus).

Immunohistochemical (IHC) staining of sections of lung was performed using a biotin-free polymer system. The paraffin-embedded sections were subjected to deparaffinization in xylene, rehydration in graded series of ethanol, and rinsed with double distilled water. Antigen retrieval was performed by immersing sections in DIVA Decloaker (Biocare Medical) at 125 °C for 30 seconds in a steam pressure decloaking chamber (Biocare Medical) followed by blocking with Background Sniper Reagent (Biocare Medical) for 10 minutes. The sections were incubated with Thyroid Transcription Factor-1 (Clone 8G7G3/1) for overnight at 4°C followed by a detection polymer system (MACH 2™; Biocare Medical). Labeled antibody was visualized by development of the chromogen (DAB Chromogen Kits; Biocare Medical).

Tissues were fixed in freshly prepared 4% paraformaldehyde for 24 h, transferred to 70% ethanol, paraffin embedded within 7-10 days, and blocks sectioned at 5 µm. Slides were baked for 30-60 min at 65°C then deparaffinized in xylene and rehydrated through a series of graded ethanol to distilled water. Heat induced epitope retrieval (HIER) was performed with the antigen retrieval buffers citraconic anhydride (0.01% with 0.05% Tween; Mx1, Iba-1), ACD P2 (ACD Cat. No. 322000; IL-6, pSTAT3), citrate buffer (pH 6.0; MPO) or 0.01M Tris-HCl pH 8.6 (pJAK2) in a Biocare NxGen Decloaking Chamber that was set to 110°C for 15 min. The slides were cooled, rinsed twice in distilled water and 1X TBS with 0.05% Tween-20 (TBS-T), blocked (TBS-T + 0.25% casein) for 30 minutes at room temperature, then incubated at room temperature with antibodies against IL-6 (Sigma; Cat. No. SAB4301665 at 1:250 for 1 hour), Mx1 (EMD; Cat. No. MABF938 at 1:1000 for 1 hour), MPO (Dako; Cat. No. A0398 at 1:1000 for 1 hour), pSTAT3 (Cell Signaling; Cat. No. 9145L at 1:200 for 2 hours), Iba-1 (BioCare; Cat. No. CP290A at 1:500 for 1 hour) and pJAK2 (abcam; Cat. No. ab32101 1:150 for 1 hour). Endogenous peroxidases were blocked with 1.5% H2O2 in TBS-T for 10 minutes. Slides were then incubated with Rabbit Polink-1 HRP (GBI Labs; Cat. No. D13-110 for MPO and Iba-1), Rabbit Polink-2 HRP (GBI Labs; Cat. No. D39-110 for IL-6, pSTAT3 and pJAK2) and Mouse Polink-2 HRP (GBI Labs; Cat. No. D37-110 for Mx1). Slides were developed using Impact™ DAB (3,3′-diaminobenzidine; Vector Laboratories), washed in ddH_2_O, counterstained with hematoxylin, mounted in Permount (Fisher Scientific), and scanned at 20x magnification on an Aperio AT2 (Leica Biosystems). Staining for MPO, Mx1, Iba-1 and pSTAT3 IHC was performed as previously described using a Biocare intelliPATH autostainer.

### Quantitative image analysis

Quantitative image analysis was performed using HALO software (v3.0.311.405; Indica Labs) on at least one lung lobe cross section from each animal. For MPO (neutrophil) and Iba-1 (macrophage) quantification, blood vessels (>5mm^2^), bronchi, bronchioles, cartilage, and connective tissue were manually excluded; subsequently, the Multiplex IHC v2.3.4 module was used to detect MPO+ or Iba-1+ cells and is presented as a proportion of total alveolar tissue (cells/mm^2^). For Mx1, the Area Quantification v2 module was used to determine the percentage of Mx1 as a proportion of the total tissue area. In all instances, manual curation was performed on each sample to ensure the annotations were accurate and to correct false positives/false negatives.

### RNAscope in situ hybridization

RNAscope in situ hybridization was performed as previously described(Chandrashekar et al., 2020a) using SARS-CoV2 anti-sense specific probe v-nCoV2019-S (ACD Cat. No. 848561) targeting the positive-sense viral RNA and SARS-CoV2 sense specific probe v-nCoV2019- orf1ab-sense (ACD Cat. No. 859151) targeting the negative-sense genomic viral RNA. In brief, after slides were deparaffinized in xylene and rehydrated through a series of graded ethanol to distilled water, retrieval was performed for 30 min in ACD P2 retrieval buffer (ACD Cat. No. 322000) at 95-98 °C, followed by treatment with protease III (ACD Cat. No. 322337) diluted 1:10 in PBS for 20 min at 40 °C. Slides were then incubated with 3% H2O2 in PBS for 10 min at room temperature. Prior to hybridization, probes stocks were centrifuged at 13,000 rpm using a microcentrifuge for 10 min, then for the antisense probe diluted 1:2 in probe diluent (ACD Cat. No. 300041) to reduce probe aggregation tissue artifacts. Slides were developed using the RNAscope® 2.5 HD Detection Reagents-RED (ACD Cat. No.322360).

### Tissue Processing

PB was collected from the femoral vein in sodium citrate, serum separation, and EDTA tubes from which plasma was separated by centrifugation within 1 hour of phlebotomy. PB was used for complete blood counts, comprehensive serum chemistry panels, and measurement of neutrophil extracellular traps (NET) activity. From EDTA PB, peripheral blood mononuclear cells (PBMCs) were isolated using a Ficoll-Paque Premium density gradient (GE Healthcare), and washed with R-10 media. R-10 media was composed of RPMI 1640 (Corning) supplemented with 10% heat-inactivated fetal bovine serum (FBS), 100 IU/mL penicillin, 100 μg/mL streptomycin, and 200 mM L-glutamine (GeminiBio). CSF was collected by inserting a 0.75-1.5-inch, 22-25-gauge needle below the external occipital protuberance into the cisterna magna and was separated by centrifugation.

Nasopharyngeal swabs were collected under anesthesia by using a clean rayon-tipped swab (ThermoFischer Scientific, BactiSwab NPG, R12300) placed approximately 2-3cm into the nares. Oropharyneal swabs were collected under anesthesia using polyester tipped swabs (Puritan Standard Polyester Tipped applicator, polystyrene handle, 25-806 2PD, VWR International) to streak the tonsils and back of throat bilaterally (throat/pharyngeal). The swabs were dipped in 1 mL viral transport media (Viral transport Media, VTM-1L, Labscoop, LLC) and vortexed for 30 sec, and the eluate was collected.

To collect BAL, a fiberoptic bronchoscope (Olympus BF-XP190 EVIS EXERA III ULTRA SLM BRNCH and BF-P190 EVIS EXERA 4.1mm) was manipulated into the trachea, directed into the primary bronchus, and secured into a distal subsegmental bronchus upon which 35-50 mL of normal saline (0.9% NaCl) was administered into the bronchus and re-aspirated to obtain a minimum of 20ml of lavage fluid. BAL was filtered through a 70μm cell strainer.

Lung tissue was cut into small pieces, using blunt end scissors, then digested using 1.5 U/mL DNase I (Roche) and 1 mg/mL of Type I collagenase (Sigma-Aldrich) using gentleMACS C tubes and gentleMACS Dissociator (miltenyi Biotec).

Hilar LN biopsies were collected at necropsy, sectioned using blunt, micro-dissection scissors and mechanically disrupted through a 70μm cell strainer and washed with R-10 media.

Mononuclear cells were counted for viability using a Countess II Automated Cell Counter (Thermo Fisher) with trypan blue stain and were cryo-preserved in aliquots of up to 2x10^7^ cells in 10% DMSO in heat-inactivated FBS. Whole tissue segments (0.5 cm^3^) were snap frozen dry, or stored in RNAlater (Qiagen), or Nuclisens lysis buffer (Biomerieux) for analyses of compound distribution, RNA-seq, and tissue viral quantification, respectively.

### Bulk and single-cell RNA-Seq Library and sequencing from NHP BALs

Single cell suspensions from BAL were prepared in BSL3 as described above for flow cytometry; for bulk RNA-Seq, 50,000 cells were lysed directly into 700 ul of QIAzol reagent. RNA was isolated using RNeasy Mini or Micro kits (Qiagen) with on-column DNase digestion. RNA quality was assessed using an Agilent Bioanalyzer and total RNA was used as input for cDNA synthesis using the Clontech SMART-Seq v4 Ultra Low Input RNA kit (Takara Bio) according to the manufacturer’s instructions. Amplified cDNA was fragmented and appended with dual-indexed bar codes using the NexteraXT DNA Library Preparation kit (Illumina). Libraries were validated by capillary electrophoresis on an Agilent 4200 TapeStation, pooled at equimolar concentrations, and sequenced on an Illumina NovaSeq6000 at 100SR, yielding 20-25 million reads per sample. For single-cell RNA-Seq, single-cell suspensions of 100,000 BAL-derived cells, and approximately 30,000 cells were loaded onto 10X Chromium Controller in the BSL3. Single cells were partitioned into droplets (Gel Beads in Emulsion: GEMs) using Chromium NextGEM Single Cell 5’ Library & Gel Bead kits on the 10X Chromium Controller(Zheng et al., 2017). The resulting cDNA was amplified and libraries were prepared for transcriptomic analysis according to manufacturer instructions. Gene expression libraries were sequenced as paired-end 26x91 reads on an Illumina NovaSeq6000 targeting a depth of 50,000 reads per cell in the Yerkes Genomics Core Laboratory(http://www.yerkes.emory.edu/nhp_genomics_core/). Cell Ranger software was used to perform demultiplexing of cellular transcript data, and mapping and annotation of UMIs and transcripts for downstream data analysis.

### Bulk RNA-Seq analysis

The quality of reads was evaluated using FastQC (https://www.bioinformatics.babraham.ac.uk/projects/fastqc/). Reads were aligned using STAR v2.7.3.(Dobin et al.). The STAR index was built by combining genome sequences for Macaca mulatta (Mmul10 Ensembl release 100), SARS-CoV2 (strain MN985325.1 - NCBI) and ERCC sequences. The gffread utility (https://github.com/gpertea/gffread) was used to convert gff3 file for SARS-CoV2 and the resulting gtf file for SARS-CoV2 was edited to include exon entries which had the same coordinates as CDS to get counts with STAR. The combined genomic and gtf files were used for generating the STAR index. Transcript abundance estimates were calculated internal to the STAR aligner using the algorithm of htseq-count(Sandler et al., 2014). The ReadsPerGene files were used to generate counts in the htseq format using a custom script that also converted the Ensembl ID to gene names using the gtf file. These files were imported in DESeq2 using the DESeqDataSetFromHTSeqCount function. DESeq2 was used for normalization(Love et al.), producing both a normalized read count table and a regularized log expression table. Only the protein coding genes defined in the gtf file were used for analysis. The design used was: ∼ Subject + Group where Group was a combination of Timepoint (baseline/2dpi/4dpi) and Condition (Untreated/Treated) factors. The regularized log expression values were obtained using the rlog function with the parameters blind =FALSE and filtType = “parametric”. The thresholds of padj < 0.05, fold-change > 1.5 and lfcSE < 1 were used to obtain significant differentially expressed genes. The VennDiagram R library was used to create the venn diagrams. GSEA 4.1.0 (https://www.gsea-msigdb.org/) was used for gene set enrichment analysis with the following gene sets: Hallmark and Canonical pathways (MsigDB), NHP ISGs (Sandler et al.) and Rheumatoid arthritis (KEGG map05323). GSEA was run with default parameters with the permutation type set to gene_set. The input for GSEA was the regularized log expression values obtained from DESeq2 which was filtered to remove genes with mean expression <=0. The regularized log expression values were also used to generate heatmaps using the Complex Heatmap R library.

### Single-cell RNA-Seq Bioinformatic Analysis

Bronchoalveolar lavage (BAL) samples from five Rhesus Macaque’s were run on 2 Nova Seq 1000 lanes and the resultant bcl files were converted to counts matrices using Cell Ranger v3.1 (10X Genomics). Further, the count matrices for each sample were processed using an inhouse single-cell RNA-seq pipeline that uses Seurat v3.0 (Satija et al., 2018) to initially integrate data from SARS-CoV-2 infected and Baricitinib treated samples. The batch corrected samples were filtered for cells expressing <250 genes, >10% Mitochondria genes, HBB, RPS and RPL genes and any doublets were removed using DoubletFinder(McGinnis et al., 2019). After filtration, the data were normalized using scTransform normalization followed by Principal Component analysis. PCs 1-30 were chosen for clustering analysis, as there was very little additional variance observed beyond PC 30. Cells were then clustered based on PC scores using the Louvian-Jaccard method. Uniform Manifold Approximation and Projection (UMAP) (McInnes et al., 2018) method was used to visualize the single cells in 2D embedding. We used Blueprint Encode database from SingleR (Aran et al., 2019) to classify cells into different cell subtypes. Differential gene expression between the clusters was assessed by MAST (Finak et al., 2015). Heatmaps, Dot plots and Violin plots were generated using ggplot2 (Wickham, 2016) package in R.

### Immunophenotyping

23-parameter flow cytometric analysis was perform on fresh PBMCs and mononuclear cells (10^6^ cells) derived from LN biopsies, BAL, and lung. Immunophenotyping was performed using anti-human monoclonal antibodies (mAbs), which we (Harper et al., 2020; McGary et al., 2017; Micci et al., 2015; Pino et al., 2019) and others, including databases maintained by the NHP Reagent Resource (MassBiologics), have shown as being cross-reactive in RMs. A panel of the following mAbs was used for longitudinal T-cell phenotyping in PBMCs: anti-CCR7-BB700 (clone 3D12; 2.5 μL; cat. # 566437), anti-CD103-BV421 (clone Ber-ACT8; 5 μL; cat. # 563882), anti-Ki-67-BV480 (clone B56; 5 μL; cat. # 566109), anti-CXCR6-BV750 (clone 13B 1E5; 2.5 μL; cat. # 747052), anti-CD3-BUV395 (clone SP34-2; 2.5 μL; cat. # 564117), anti-CD8-BUV496 (clone RPA-T8; 2.5 μL; cat. # 612942), anti-CD45-BUV563 (clone D058-1283; 2.5 μL; cat. # 741414), anti-CD49a-BUV661 (clone SR84; 2.5 μL; cat. # 750628), anti-CD28-BUV737 (clone CD28.2; 5 μL; cat. # 612815), anti-CD69-BUV805 (clone FN50; 2.5 μL; cat. # 748763), and Fixable Viability Stain 700 (2 μL; cat. # 564997) all from BD Biosciences; anti-CD95-BV605 (clone DX2; 5 μL; cat. # 305628), anti-HLA-DR-BV650 (clone L243; 5 μL; cat. # 307650), anti-CD25-BV711 (clone BC96; 5 μL; cat. # 302636), anti-PD-1-BV785 (clone EH12.2H7; 5 μL; cat. # 329930), anti-CD101-PE-Cy7 (clone BB27; 2.5 μL; cat. # 331014), anti-FoxP3-AF647 (clone 150D; 5 μL; cat. # 320014), and anti-CD4-APC-Cy7 (clone OKT4; 2.5 μL; cat. # 317418) all from Biolegend; anti-CD38-FITC (clone AT1; 5 μL; cat. # 60131FI) from STEMCELL Technologies; and anti-CXCR5-PE (clone MU5UBEE; 5 μL; cat. # 12-9185-42), anti-GranzymeB-PE-TexasRed (clone GB11; 2.5 μL; cat. # GRB17), and anti-CD127-PE-Cy5 (clone eBioRDR5; 5 μL; cat. # 15-1278-42) all from Thermo Fisher (**Supplementary Fig. 4**). mAbs for chemokine receptors (i.e. CCR7) were incubate at 37°C for 15 min, and cells were fixed and permeabilized for 30 min at room temperature using a FoxP3 / Transcription Factor Staining Buffer Kit (Tonbo Biosciences; cat. # TNB-0607-KIT). A panel of the following mAbs was used for the longitudinal phenotyping of innate immune cells in whole blood (500 μL), as described in (Kuri-Cervantes et al., 2020), and mononuclear cells (10^6^ cells) derived from LN biopsies, BAL, and lung: anti-CD20-BB700 (clone 2H7; 2.5 μL; cat. # 745889), anti-Ki-67-BV480 (clone B56; 5 μL; cat. # 566109), anti-CD14-BV605 (clone M5E2; 2.5 μL; cat. # 564054), anti-CD56-BV711 (clone B159; 2.5 μL; cat. # 740781), anti-CD115-BV750 (clone 9-4D2-1E4; 2.5 μL; cat. # 747093), anti-CD3-BUV395 (clone SP34-2; 2.5 μL; cat. # 564117), anti-CD8-BUV496 (clone RPA-T8; 2.5 μL; cat. # 612942), anti-CD45-BUV563 (clone D058-1283; 2.5 μL; cat. # 741414), anti-CCR2-BUV661 (clone LS132.1D9; 2.5 μL; cat. # 750472), anti-CD16-UV737 (clone 3G8; 2.5 μL; cat. # 564434), anti-CD69-BUV805 (clone FN50; 2.5 μL; cat. # 748763), and Fixable Viability Stain 700 (2 μL; cat. # 564997) all from BD Biosciences; anti-CD38-FITC (clone AT1; 2.5 μL; cat. # 60131FI) from STEMCELL Technologies; anti-CD161-BV421 (clone HP-3G10; 5 μL; cat. # 339914), anti-HLA-DR-BV650 (clone L243; 5 μL; cat. # 307650), anti-CD11c-BV785 (clone 3.9; 5 μL; cat. # 301644), anti-CD11b-PE (clone ICRF44; 2.5 μL; cat. # 301306), and anti-CD123-APC-Fire750 (clone 315; 2.5 μL; cat. # 306042) all from Biolegend; anti-GranzymeB-PE-TexasRed (clone GB11; 2.5 μL; cat. # GRB17) from Thermo Fisher; anti-CD66abce-PE-Vio770 (clone TET2; 1 μL; cat. # 130-119-849) from Miltenyi Biotec; and anti-CD27-PE-Cy5 (clone 1A4CD27; 2.5 μL; cat. # 6607107) and anti-NKG2A-APC (clone Z199; 5 μL; cat. # A60797) from Beckman Coulter (**Supplementary Fig. 4**). mAbs for chemokine receptors (i.e. CCR2) were incubated at 37°C for 15 min, and cells were fixed and permeabilized at room temperature for 15 min with Fixation/Permeabilization Solution Kit (BD Biosciences; cat. #554714). For each sample a minimum of 1.2x10^5^ stopping gate events (live CD3^+^ T-cells) were recorded except for RB in which a minimum of 5x10^4^ stopping gate events were recorded. All samples were fixed with 4% paraformaldehyde and acquired within 24 hours of fixation. Acquisition of data was performed on a FACSymphony A5 (BD Biosciences) driven by FACS DiVa software and analyzed with FlowJo (version 10.7; Becton, Dickinson, and Company).

Single cells were then selected using FSC-A x FCS-H gate. A lymphocyte and granulocyte gate based on FSC-A and SSC-A was defined. Live cells were gated followed by CD45^+^ cells. UMAP analysis (Uniform Manifold Approximation and Projection for Dimension Reduction) was performed in live CD45^+^for unbiased evaluation of the distribution of the key markers. Projection of the density of cells expressing markers of interest were visualized/plotted on a 2-dimensional UMAP (https://arxiv.org/abs/1802.03426, https://github.com/lmcinnes/umap). We used the Phenograph clustering approach (https://github.com/jacoblevine/PhenoGraph)

### Determination of intracellular cytokine induction following SARS-CoV-2 S peptide pool and PMA/Ionomycin stimulation

Cryo-preserved PBMCs were thawed, resuspended in RPMI medium supplemented to contain a final concentration of 10% Fetal Bovine Serum (FBS) (Corning Life Sciences/Media Tech Inc, Manassas, VA), 10mM HEPES, 1x MEM nonessential amino acids (Corning Life Sciences/Media Tech Inc, Manassas, VA), 1mM Sodium Pyruvate (Lonza, Walkersville, MD, U.S.A), 1mM Penicillin/Streptomycin containing Amphothericin B (Sigma Life Sciences, St Louis, MO, U.S.A) and 1x 2-Mercaptoethanol (GIBCO, Invitrogen, Carlsbad, CA, U.S.A). PBMCs were rested overnight at 37°C in a cell culture incubator. Cells were then stimulated for detection of cytokine production by T cells as described before (Kasturi et al., 2020). Briefly, 2 x 10^6^ cells were cultured in 200μL final volume in 5mL polypropylene tubes (BD Biosciences, San Diego, CA, U.S.A) in the presence of anti-CD28 (1μg/mL) and anti-CD49d (1μg/mL) [BD Biosciences] and the following conditions; a) negative control with DMSO only, b) S peptide pool (Grifoni et al., 2020) and c) PMA/Ionomycin in the presence of Golgi transport inhibitors - 10 μg/mL of Brefeldin A (Sigma-Aldrich). After stimulation, cells were washed and stained for cell surface antigens with two panels. Panel 1: anti-CD3 BUV395 (clone SP34-2; 2.5 μL; cat. # 564117), anti-CD8-BUV496 (clone RPA-T8; 2.5 μL; cat. # 612942), and Fixable Viability Stain 700 (2 μL; cat. # 564997) all from BD Bioscience; anti-CD4 APC/Cy7 (clone OKT4; 2.5 μL; cat. # 317418) from Biolegend; To detect intracellular expression of cytokines, mononuclear cells were fixed and permeabilized with a Cytofix/Cytoperm kit (BD Biosciences) and stained as follows: anti-IL-21 BV421 (clone 3A3-N2.1; 5 μL; cat# 564755) from BD Bioscience; anti-IL-2 BV650 (clone MQ1-17H12; 5 μL; cat# 500334) and anti-IFNγ PE/Dazzle 594 (clone B27; 5 μL; cat# 506530) both from Biolegend; anti-IL-17a Alexa Fluor 488 (clone eBio64DEC17; 5 μL; cat# 53-7179-42), anti-IL-22 APC (clone IL22JOP; 5 μL; cat# 17-7222-82), and anti-TNFα PE-Cyanine7 (clone Mab11; 0.5 μL; cat# 25-7349-82) all from Thermo Fisher Scientific; and anti-IL-4 PE (clone 7A3-3; 5 μL; cat# 130-091-647) from Miltenyl Biotech. Panel 2: anti-IL-2 Alexa Fluor 488 (clone MQ1-17H12; 3 μL; cat# 500314), anti-CD8a PerCP Cy5.5 (clone RPA-T8; 3 μL; cat# 301032), anti-CD4 BV421 (clone OKT4; 2.5 μL; cat# 317434), and anti-IFNγ Alexa 647 (clone 4S.B3; 3 μL; cat# 502516) from Biolegend; anti-CD3 BV605 (clone SP34-2; 2 μL; cat# 562994) from BD Biosciences; anti-IL-4 PE (clone 7A3-3; 5 μL; cat# 130-091-647) from Miltenyl Biotech; anti-IL-17a PE-efluor 610 (clone eBio64DEC17; 3 μL; cat# 61-7179-42), anti-TNFα PE-Cyanine7 (clone Mab11; 0.5 μL; cat# 25-7349-82), and Live Dead APC-Cy7 (1:1000; cat# 65086514) from Thermo Fisher Scientific. The frequency of SARS-CoV-2 specific CD4^+^ and CD8^+^ T-cells producing single or multiple cytokines was determined after background subtraction. All samples were fixed with 4% paraformaldehyde and acquired within 24 hours of fixation. Acquisition of data was performed on a FACSymphony A5 (BD Biosciences) driven by FACS DiVa software and analyzed with FlowJo (version 10.7; Becton, Dickinson, and Company).

### Isolation of Non-human Primate Neutrophils

Neutrophils were obtained from peripheral blood of SARS-Cov-2 infected Rhesus Macaques 5 days pre-infection and at days 4, 7, and 10 post-infection. Peripheral blood (0.5-1 ml) was collected using a citrate containing Vacutainer and the upper serum layer was removed. The red blood cell layer was lysed with 2 mL of Red Blood Cell Lysis Buffer (Cat# 11814389001, Roche) in a 15 mL tube. The tube was gently inverted for 10 minutes at room temperature and centrifuged at 500 x *g* for 7 minutes at room temperature. This step was repeated gently inverting for 5 minutes. Following centrifugation, the cell pellet was re-suspended in a final volume of 2 mL of 1x PBS/EDTA buffer gently. Cells were centrifuged at 500 x *g* for 7 mins a room temperature and the leukocyte pellet was re-suspended in 1 mL 1x PBS/EDTA buffer and carefully overlaid onto 3 mL of 65% Percoll/EDTA solution. The Percoll cell gradient was centrifuged at 400 x g for 20 mins at room temperature with the brake turned off. The neutrophil cell layer was collected, re-suspended/washed with 5 mL of 1x PBS/EDTA buffer and centrifuged at 500 x g for 10 minutes at room temperature. The neutrophil cellular pellet was re-suspended in RPMI 1640 media. Purification of the cell fragment was confirmed using flow cytometry and Wright Giemsa staining.

### Quantification of extracellular DNA using SYTOX green Assay

Abundance of extracellular DNA, a surrogate of NETs, was quantified using the SYTOX green assay. Freshly, isolated non-human primate neutrophils were plated onto a 96-well plate at a density of 10^5^ cells per well in 100 μL RPMI 1640 media then stimulated with 50 μg/mL LPS to induce NET formation. SYTOX green dye (5 μM, #S7020; Invitrogen, Carlsbad, CA) was added to each well and the fluorescence intensity was read with a filter setting at 485-nm excitation/525- nm emission using a Synergy H1 Microplate Reader and Gene5 software (Biotek, Winooski, VT). A fluorescence reading was collected every 15 mins for a total of 2 hours at 37°C. Images of the fluorescent cells were immediately taken using a fluorescent microscope (Olympus).

### Statistical Analyses

All statistical analyses were performed two-sided with p-values ≤0.05 deemed significant. Ranges of significance were graphically annotated as follows: *, p<0.05; **, p<0.01; ***, p<0.001; ****, p<0.0001. Due to the low number of animals included in our study, p values ≤0.1 have been indicated in the graphs. Analyses, unless otherwise noted, were performed with Prism version 8 (GraphPad).

### Data Availability Statement

Source data supporting this work are available from the corresponding author upon reasonable request. The following sequencing data have been deposited in GenBank: SARS-CoV-2 viral stock (accession # PENDING). Data tables for expression counts for bulk and single-cell RNA-Seq for BAL are deposited in NCBI’s Gene Expression Omnibus and are accessible through GEO accession #####. Custom scripts and supporting documentation on the RNA-Seq analyses will be made available at https://github.com/BosingerLab/.

---

**Figure S1.** (**a - left**) Concentration of baricitinib 2 hours post-dosing in brain and CSF and (**a - right**) 24 hours post-dosing in CSF.

**Figure S2.** Circulating immune cell populations, and blood chemistry parameters in SARS-CoV-2 infected RMs. Longitudinal frequency of (**a)** monocyte, (**b)** neutrophils, and (**c)** lymphocytes in blood of SARS-CoV-2 infected RMs. (**d)** Red blood cell counts (RBC), (**e)** hematocrit (HCT), (**f)** hemoglobin (HGB) and (**g)** alkalin phosphatase (ALP) levels were analyzed throughout the study. Different symbols represent individual animals. Bold lines represent the average of the baricitinib treated group (blue), and the untreated group (red).

**Figure S3.** Representative images of *in situ* RNA hybridization (RNAscope) targeting viral RNA strands identifying clusters of infected cells within the lung parenchyma in both treated and untreated SARS-CoV-2 infected animals. Scale bars: 500um.

**Figure S4.** Representative immunohistochemistry (IHC) images of (**a**) neutrophils (myeloperoxidase+, MPO, cells) (**c**) macrophages (ionized calcium-binding adaptor molecule 1+, Iba-1, cells), and (**e**) Interferon-induced GTP-binding protein+ (Mx1) cells in lungs of baricitinib treated and untreated SARS-CoV-2 infected RMs. Scale bars 500 μm. Quantification of (**b**) neutrophils (MPO+ positive cells/mm^2^), (**d**) macrophages (lba-1+ cells/mm^2^), and (**f**) Interferon-induced protein Mx1 (% area total lung Mx1+) in IHC lung images of baricitinib treated, and untreated controls of SARS-CoV-2 infected RMs, and uninfected RMs.

**Figure S5.** Cross-sectional GSEA analysis comparing 4 DPI untreated vs 4 DPI baricitinib treated, or 2 DPI untreated vs 2 DPI baricitinib treated. (**a-c**) GSEA comparisons of 4 DPI untreated vs 4 DPI baricitinib treated are shown as black symbols, and comparisons of or 2 DPI untreated vs 2 DPI baricitinib treated are shown as grey symbols. (**a**) GSEA enrichment plots for the GSEA line plot of HALLMARK_IL6_JAK_STAT3_SIGNALING pathway (MSIGDB# M5897). (**b**) GSEA line plot of HALLMARK_TNFA_SIGNALING_VIA_NFKB pathway (MSIGDB # M5890). (**c**) GSEA line plot of REACTOME NEUTROPHIL DEGRANULATION gene set (REACTOME #M27620). (**d**) Heatmap of leading edge genes for REACTOME NEUTROPHIL DEGRANULATION gene set based on untreated 4 DPI vs baseline contrast. The log2 expression and the reference is the median of all baseline samples as indicated at right. The top 35 genes are shown in order of GSEA analysis of the cross-sectional 4 DPI comparison.

**Figure S6.** GSEA analysis for KEGG Rheumatoid Arthritis gene set (**a**) GSEA contrasting 4 DPI vs 2 DPI for untreated and treated arms. GSEA curves are colored by experimental arm. Leading edge genes are indicated by solid dots. The hash plot under GSEA curves indicate individual genes and their rank in the dataset. Left-leaning curves (i.e. positive enrichment scores) indicate enrichment at 4 DPI, right-leaning curves (negative enrichment scores) indicate higher enrichment at 2 DPI, and sigmoidal curves indicate a lack of enrichment, i.e. equivalent expression between the groups being compared. The normalized enrichment scores and nominal p-values testing the significance of each comparison are indicated. (**b**) GSEA comparisons of 4 DPI untreated vs 4 DPI baricitinib treated samples (black symbols); comparisons of 2 DPI untreated vs 2 DPI baricitinib treated samples (grey symbols). (**c**) plot showing log10 average normalized counts obtained from DESeq2 for leading edge genes at 2 DPI in untreated and treated samples, and at 4 DPI.

**Figure S7.** (**a**) Expression as UMAP projection of interferon stimulated genes in macrophages for treated and untreated samples at baseline and 4 DPI. (**b**) Heatmap showing average expression of genes of interest in macrophages for treated and untreated samples at baseline and 4 DPI. (**c-e**) Dot plots representing gene expression levels and percentage of cells expressing genes associated with inflammation, chemokine response and interferon stimulation

**Figure S8.** (**a-c**) Expression as UMAP projection of inflammation, chemokine and interferon stimulated genes across major cell types in BAL for treated and untreated samples at baseline and 4 DPI.

**Figure S9.** Representative gating strategy of (a) neutrophils, and (b) T cell populations analyzed in the study.

**Figure S10.** Frequency of circulating CD4+ T cells spontaneously (without stimulation) producing pro-inflammatory Th17 related cytokines (IL-17^+^; IL-17^+^IL-21^+^; IL-17^+^IL-22^+^) at necropsy (days 10-11 post infection) in baricitinib (blue) and untreated (red) SARS-CoV-2 infected RMs. Different symbols indicate different animals. Bars represent the average values per group.

**Figure S11. Treatment with baricitinib does not affect the immune T cell responses in SARS-CoV-2 infected RMs.** (**a**) Representative flow cytometry staining of IFNγ, TNFα, IL-2, IL-4 and IL-17a in CD4^+^ and CD8^+^ T cells of a SARS-CoV-2 infected RM following stimulation with SARS-CoV-2 S peptide pool. IFNγ, Unstimulated background values were subtracted from S peptide stimulated values to determine T cell cytokine. TNFα, IL-2, IL-4 and IL-17a frequency levels in (**b**) CD4^+^ and (**c**) CD8^+^T cells following stimulation with SARS-CoV-2 S peptide pool. IFNγ, TNFα, IL-2, IL-4 and IL-17a frequency levels in (**d**) CD4^+^ and (**e**) CD8^+^T cells following stimulation with PMA/Ionomycin. Values from unstimulated controls were subtracted in all cases. Granzyme B and PD-1 levels in (**f and g**) blood and (**h and i**) BAL memory CD8^+^T cells measured by flow cytometry. Each symbol represents individual animals. Thick lines represent the average of the baricitinib treated (blue line), and untreated groups (red line). Bars represent the average of the treated and untreated groups. Statistical analysis was performed using a non-parametric Mann-Whitney Test.

