## Supplementary figures and images for "Baricitinib treatment resolves lower airway inflammation and neutrophil recruitment in SARS-CoV-2-infected rhesus macaques"

### Figure S1

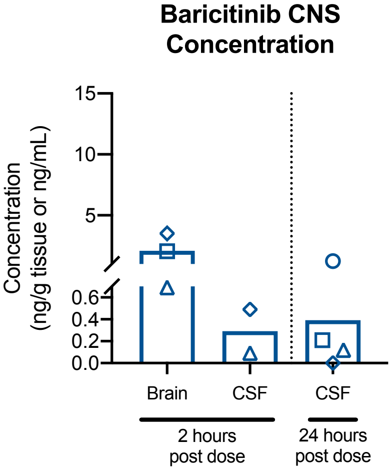

### Figure S2

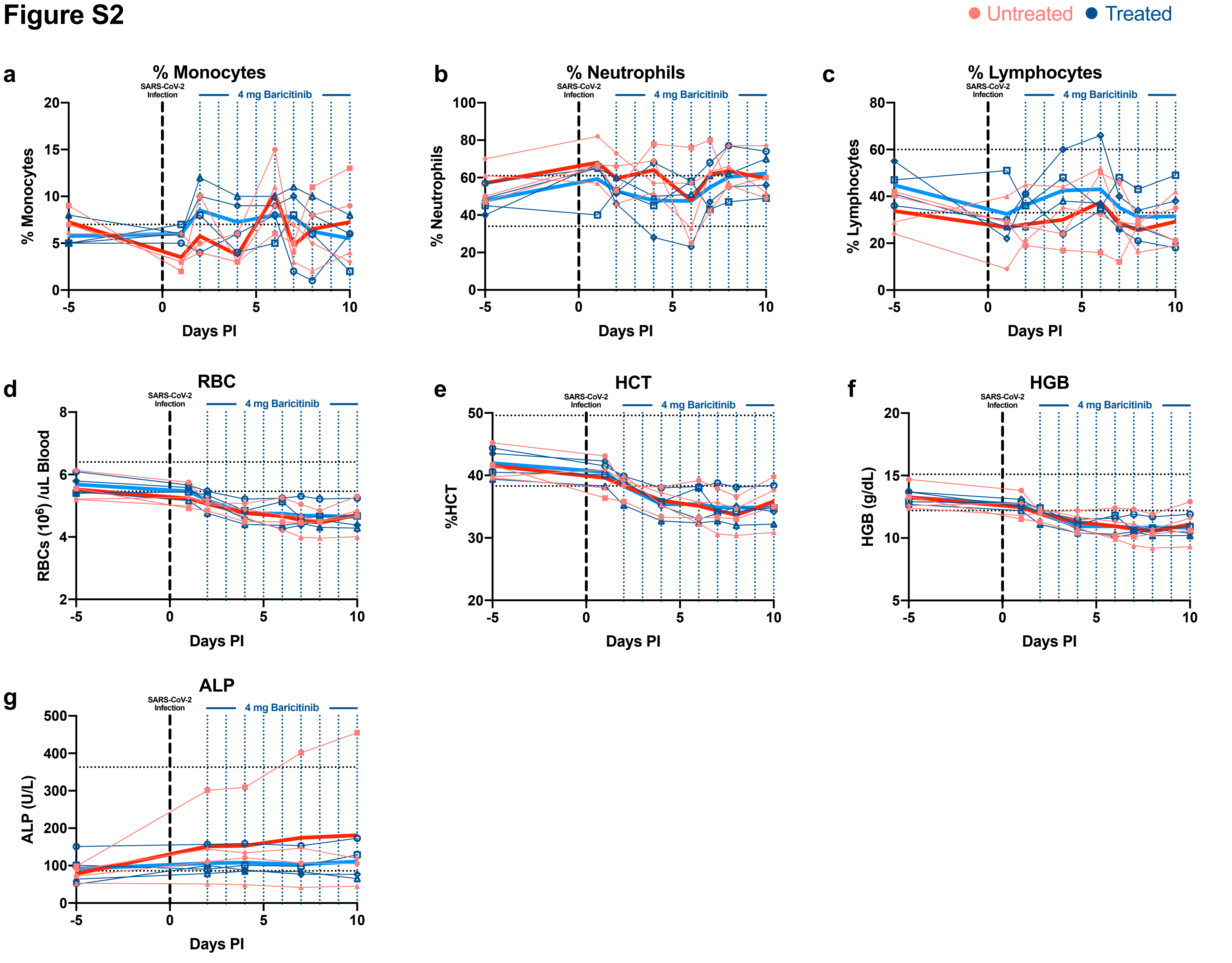

### Figure S3

#### Figure S3

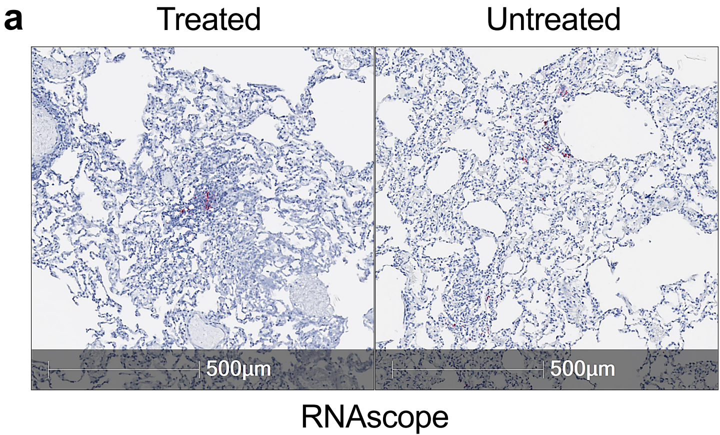

### Figure S4

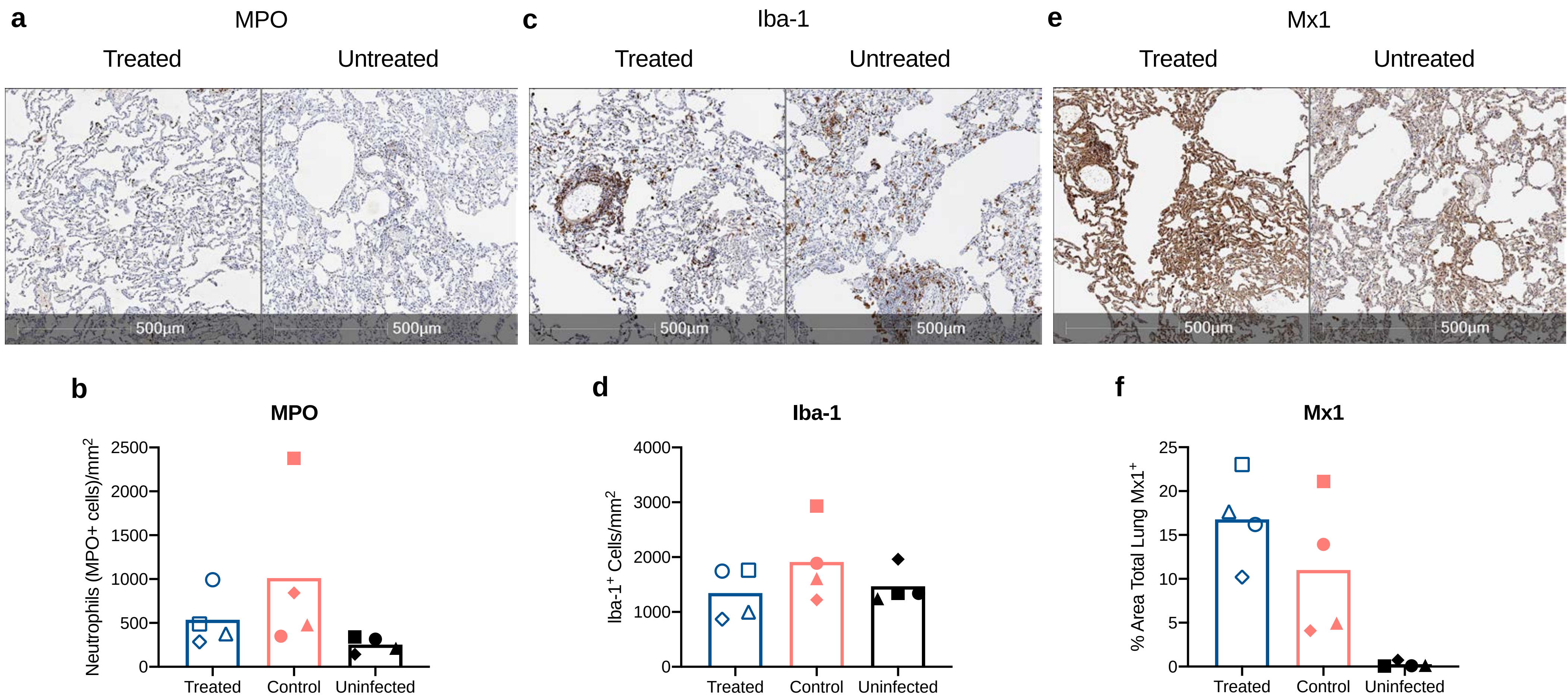

### Figure S5

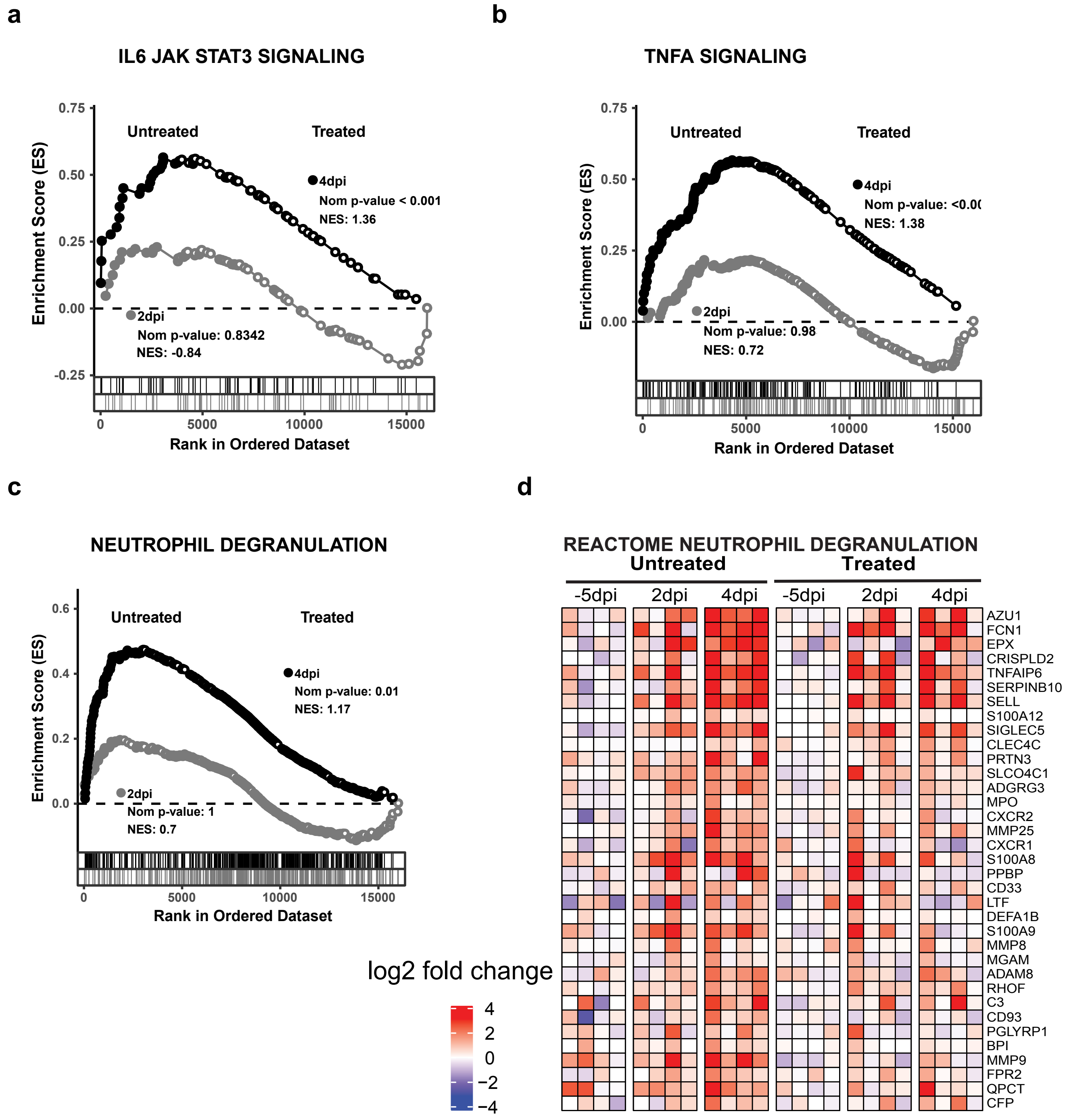

### Figure S6

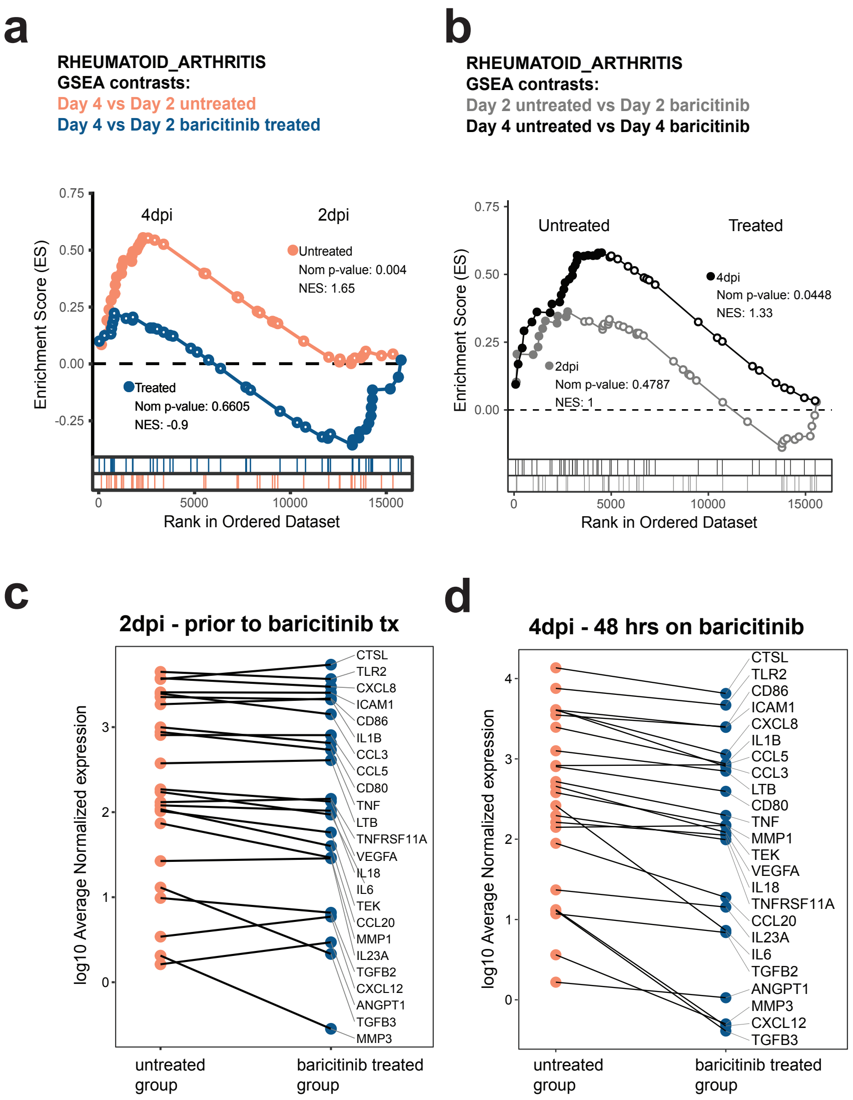

### Figure S7

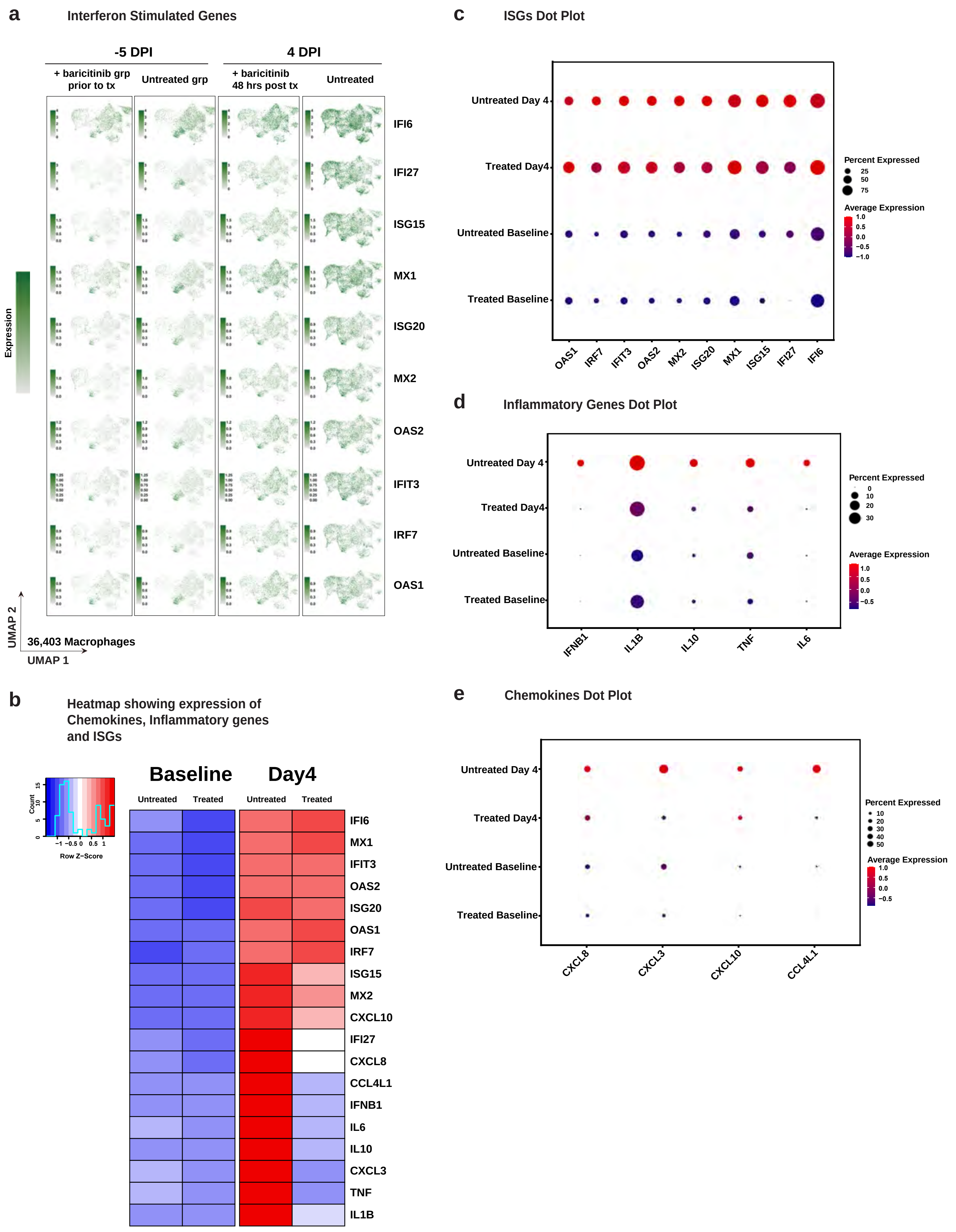

### Figure S8

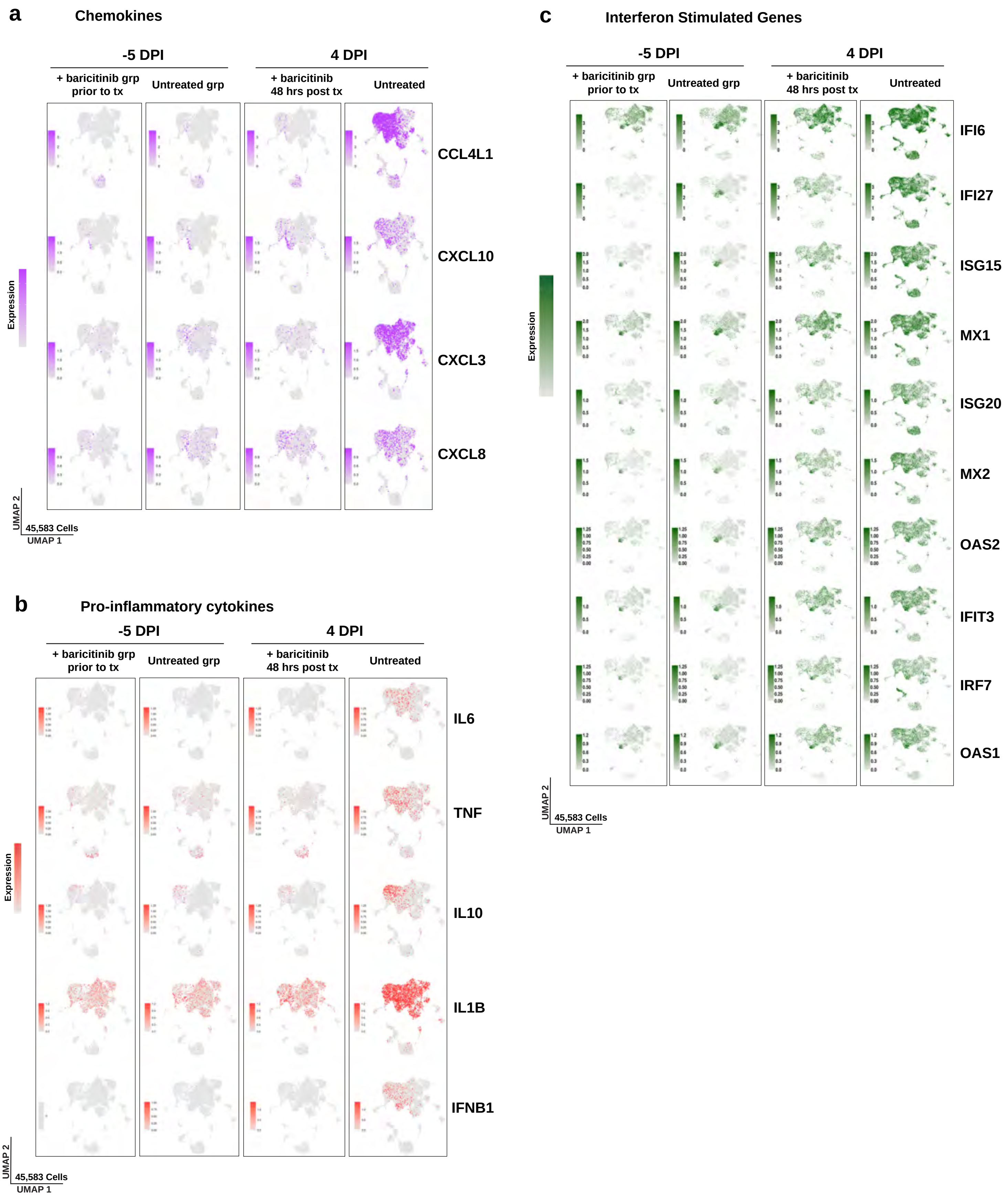

### Figure S9

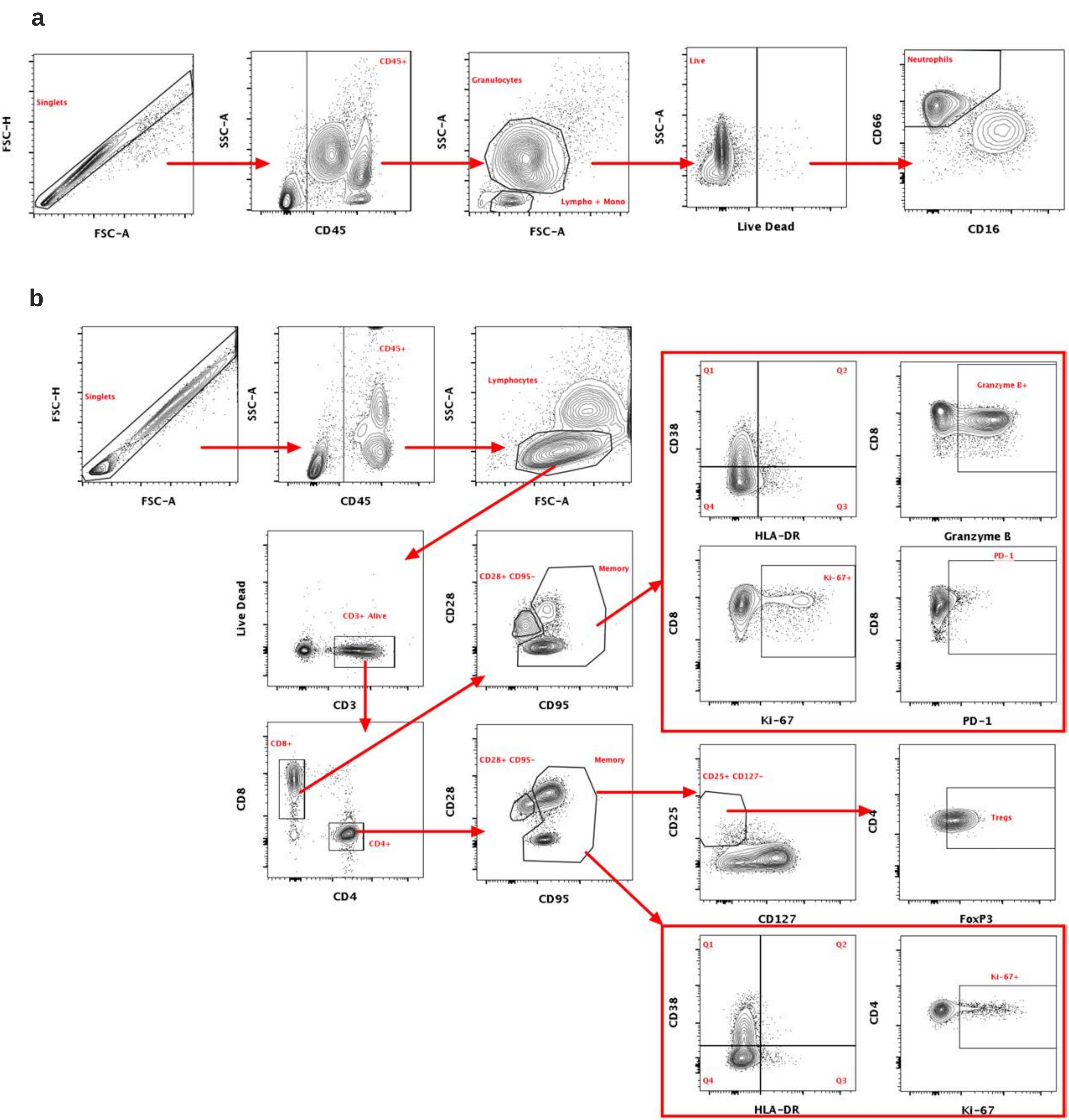

### Figure S10

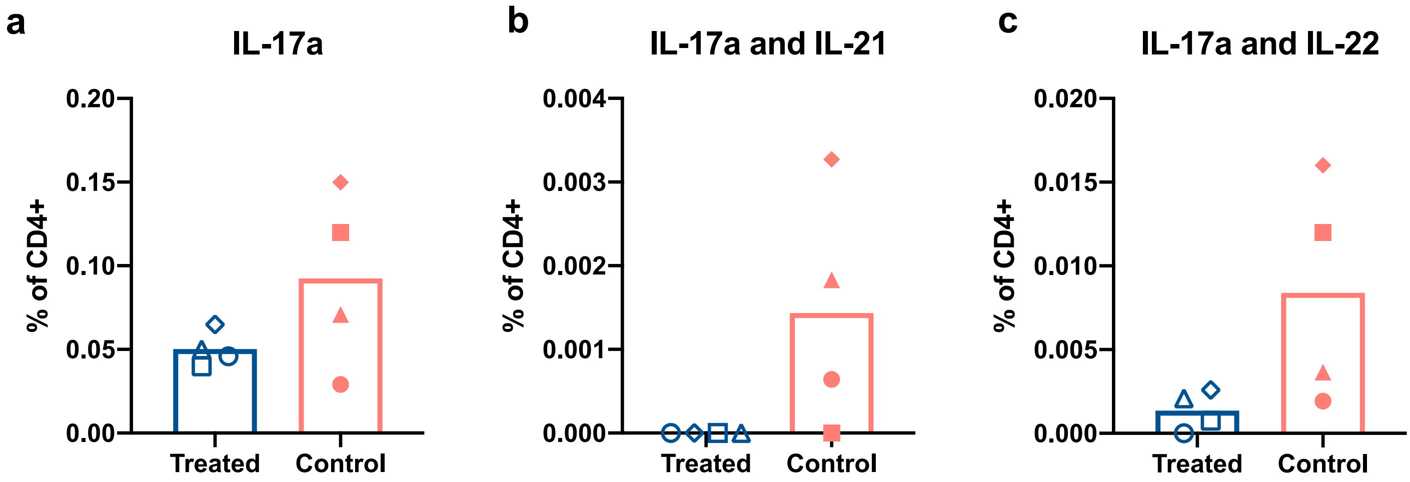

### Figure S11

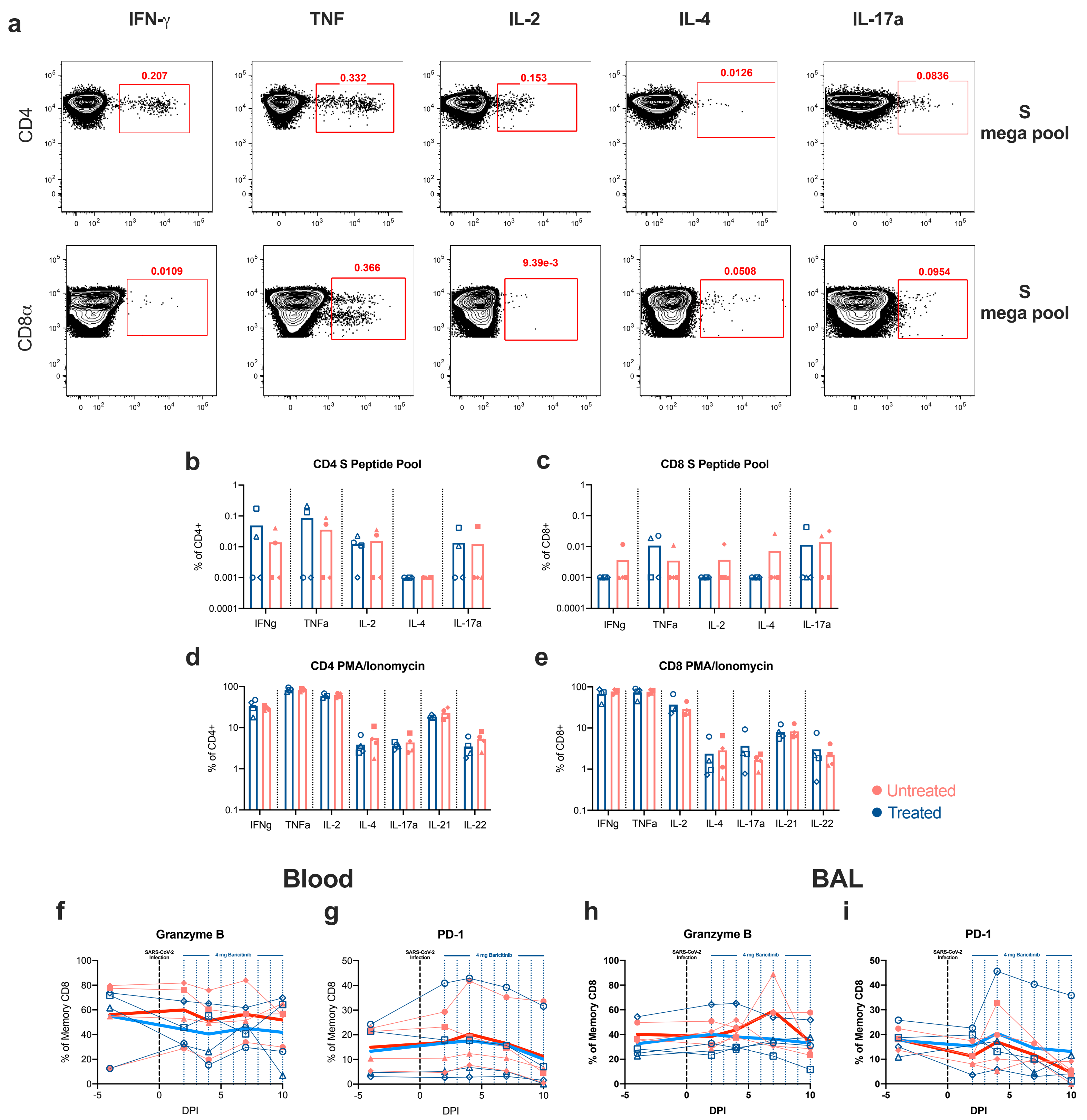
